# Role for DNA double strand end-resection activity of RecBCD in control of aberrant chromosomal replication initiation in *Escherichia coli*

**DOI:** 10.1101/2022.05.28.493860

**Authors:** Sayantan Goswami, Jayaraman Gowrishankar

## Abstract

Replication of the circular bacterial chromosome is initiated from a locus *oriC* with the aid of an essential protein DnaA. One approach to identify factors acting to prevent aberrant *oriC-*independent replication initiation in *Escherichia coli* has been that to obtain mutants which survive loss of DnaA. Here we show that a Δ*recD* mutation, associated with attenuation of RecBCD’s DNA double strand end-resection activity, provokes abnormal replication and rescues Δ*dnaA* lethality in two situations: (i) in absence of 5’-3’ single-strand DNA exonuclease RecJ, or (ii) when multiple two-ended DNA double strand breaks (DSBs) are generated either by I-SceI endonucleolytic cleavages or by radiomimetic agents phleomycin or bleomycin. One-ended DSBs in the Δ*recD* mutant did not rescue Δ*dnaA* lethality. With two-ended DSBs in the Δ*recD* strain, Δ*dnaA* viability was retained even after linearization of the chromosome. Data from genome-wide DNA copy number determinations in Δ*dnaA*-rescued cells lead us to propose a model that nuclease-mediated DNA resection activity of RecBCD is critical for prevention of a σ-mode of rolling-circle over-replication when convergent replication forks merge and fuse, as may be expected to occur during normal replication at the chromosomal terminus region or during repair of two-ended DSBs following “ends-in” replication.

## Introduction

A requirement for timely and faithful duplication of a cell’s genetic material is an essential feature of all life forms (1). The genome of most eubacteria, of which *Escherichia coli* is the prototype, is comprised of a single circular chromosome. Initiation of chromosomal replication is exquisitely controlled as to occur once per cell division from a single locus *oriC*, with the aid of an essential initiator protein DnaA (1–3); concomitant protein synthesis is necessary for fresh rounds of initiation from *oriC*, although the reasons for this requirement are not fully understood (4).

From *oriC*, replication then proceeds bidirectionally across the pair of “replichores” for the forks to then meet and merge within a broad terminus region that is situated diametrically opposite *oriC* [reviewed in (5, 6)]. The terminus region is flanked at its ends by *Ter* sequences which represent binding sites for the Tus protein; the Tus-bound *Ter* sites act in a polar fashion to block progression of a replisome that has already traversed the terminus region from entering the opposite replichore, thus serving to limit most if not all fork mergers to occur within this region [reviewed in (5, 7, 8)]. Additional *Ter* sequences are also present on either replichore in the *oriC-* distal half of the chromosome, each so oriented as to block replisome progression towards *oriC* (5, 7, 8).

Replisomes often encounter impediments to their progression (in the form of lesions in, or protein complexes bound to, DNA) which may lead to their pausing, collapse or disintegration, and thereby to generation of a single broken DNA end or “one-ended” double strand break (DSB) (9–11). Mechanisms of repair by homologous recombination and of replication restart then operate at each of these sites to reconstitute a fresh replisome that resumes its directional progression towards the terminus [reviewed in (11, 12)]. Recombinational repair is comprised of the steps of (i) DNA resection from a double-strand (ds) DNA end that is mediated by the RecBCD complex (which possesses DNA helicase and nuclease activities), followed by (ii) RecA-mediated synapsis of a single-strand (ss) DNA 3’ end into a homologous DNA duplex to generate a D-loop [reviewed in (12–18)]. Replication restart occurs from the D-loop and is mediated by the PriABC and DnaT proteins through one or more redundant pathways (9, 19, 20). RecA’s binding to ss-DNA also triggers an “SOS response”, during which the LexA repressor is cleaved so that expression of the LexA (also called SOS) regulon is transcriptionally upregulated to aid in cell survival following DNA damage (21).

Incidentally, the same mechanisms of recombinational repair and replication restart are presumed also to operate at sites of two-ended DSBs to mediate “ends-in” replication and thus to restore circularity of the chromosome (9, 14, 16, 19). Generation of such two-ended DSBs is replication-unrelated, and occurs upon the action of endonucleases such as I-SceI or upon exposure to ionizing radiation or radiomimetic agents (22, 23) like phleomycin (Phleo) or bleomycin (Bleo).

The organization and replication of the chromosome of *E. coli* is characterized by several additional (and probably inter-dependent) features of interest, all of which are correlated with the *ori-*to-*Ter* directionality of its two replichores. First, the majority of heavily transcribed genes (including all seven ribosomal RNA [*rrn*] operons) are codirectional with replisome progression (5), and there is evidence that head-on transcription-replication conflicts engendered either by inversion of *rrn* operons (24) or by ectopic placement of *oriC* (25) is detrimental. Second, the chromosome is divided into a number of macrodomains that are symmetrically distributed across the two replichores (26, 27), although the details of such organization and their contribution to bacterial fitness are still under investigation. Third, the mechanisms of decatenation and segregation of sister chromatids, as well as those of cell division, have evolved to be coupled to replication of the chromosome terminus region (6).

A fourth feature of interest is the bias in distribution of two sets of *cis* sequences (Chi and KOPS) across the genome, that are both involved in functions related to the resolution of problems arising from collapsed or disintegrated replication forks (18, 28). The distributional bias of Chi serves to limit the extent of RecBCD-mediated DNA resection at one-ended DSB sites (13, 18), whereas that of KOPS is intended to facilitate monomerization of chromosome dimers that may be generated following recombinational repair (28).

Finally, *ori-*to-*Ter* directionality of replisome progression also imposes a defined gene dosage gradient across the two replichores in asynchronously growing populations of cells (5, 29). The steepness of this gradient is accentuated in rapidly growing cultures because of the phenomenon of “multi-fork replication”, wherein new rounds of replication are initiated at *oriC* even as the earlier pairs of forks have not reached the terminus region (29, 30). Accordingly, highly expressed genes have also been favoured in evolution to be located closer to *oriC* than to the terminus (31).

Given that all of the features above require that chromosomal replication be initiated from the singular *oriC* locus, it is not surprising that mechanisms also exist in *E. coli* cells for avoidance of aberrant *oriC*-independent chromosomal replication. Such *oriC-*independent replication is also called stable DNA replication (SDR) since it persists even after transcription or translation have been inhibited in the cells (4).

One approach to delineate the mechanisms for avoidance of aberrant replication has been that to obtain mutants which retain viability in absence of DnaA (whose action is obligatory for replication initiation from *oriC*). Such mutants are said to exhibit constitutive SDR (4)]. The mutations that have been so identified include those abrogating the functions of RNase HI (4, 32, 33); RecG helicase (32, 34); topoisomerase I (35); or Dam DNA methylase (36). Rescue of lethality associated with DnaA deficiency also occurs upon loss of different combinations of DNA nucleases Exo I, Exo VII, SbcCD, and RecJ (37). Of these, the last is a 5’ ss-DNA exonuclease (38), whereas the first three have been reported to function redundantly as 3’ ss-DNA exonucleases (37); however, Exo VII also possesses 5’ ss-DNA exonuclease activity (38, 39), and SbcCD cleaves hairpin DNA and also degrades duplex DNA by a complex catalytic mechanism (40–42).

In all of the cases above, two additional mutations (Δ*tus* and *rpoB*35*) are also required for (or, in case of RNase HI deficiency, facilitate) DnaA-independent viability (32, 34–37); the two mutations presumably aid in the progression of *oriC-*independent replication around the circular chromosome, the former by disrupting the polar replication barriers at *Ter* sequences (7, 8) and the latter by resolving head-on collisions of transcription complexes with the replisome (43–46).

To further explore the mechanisms for avoidance of SDR, we have examined additional conditions that serve to confer Δ*dnaA* viability in *E. coli*. Our results suggest that ds-DNA end-resection activity of RecBCD [which is attenuated in Δ*recD* mutants (12, 13)] is critically required to prevent a rolling-circle type (that is, σ-mode) of over-replication from sites where convergent replication forks merge and fuse. Such fork mergers are expected to occur both during normal replication in the chromosomal terminus region, as well as during “ends-in” replication for repair of two-ended DSBs.

## Materials and Methods

### Bacterial strains and plasmids

Genotypes of *E. coli* strains are listed in Table S1. Knockout (Kan^R^ insertion-deletion) alleles sourced from the Keio collection (47) included the following: *dinG, rdgB, recA, recD, recG, recJ, sbcC, tus, xseA*, and *xonA*. The construction by recombineering of strains with distributed I-SceI sites on the chromosome is described in the *Supplementary Text*.

Previously described plasmids include: pKD13, pKD46, and pCP20, for use in recombineering experiments and for Flp-mediated site-specific excision of FRT-flanked DNA segments (48); ASKA vector pCA24N and its derivatives expressing *recD*^*+*^ or *recJ*^*+*^ (49); and the shelter plasmids pHYD2388 and pHYD4805 each carrying *dnaA*^*+*^ (36) and pHYD5701 carrying *recA*^*+*^ (36). Construction of plasmid pHYD5301 (used for recombineering of I-SceI sites at different chromosomal locations) is described in the *Supplementary Text*.

### Culture media and conditions

Strains were routinely cultured in LB medium at 37° (50), and supplementation of growth media with Xgal or antibiotics ampicillin (Amp), chloramphenicol (Cm), kanamycin (Kan), spectinomycin (Sp), tetracycline (Tet), and trimethoprim (Tp) were at the concentrations described earlier (51). Phleo and Bleo were each added at 1 μg/ml, and L-arabinose (Ara) at 0.2%.

### Blue-white assay for determining viability of Δ*dnaA* and Δ*recA* derivatives

The ‘blue-white screening’ strategy used to identify conditions that render a Δ*dnaA* mutant viable was as earlier described (35, 36). The parental strain is Δ*dnaA* (and Δ*lacZ*) on the chromosome and carries a single-copy-number Tp^R^ *dnaA*^*+*^ *lacZ*^*+*^ shelter plasmid pHYD2388 whose equi-partitioning to daughter cells during cell division is inefficient; thus, typically around 5 to 20% of cells in a culture (grown in absence of Tp) are plasmid-free. The latter cells would be able to form colonies (white on Xgal-supplemented medium) only if SDR in them was sufficient to confer viability. [Since plasmid loss frequency in this assay is relatively low, occurrence of white-sectored blue colonies is not as common as that in an analogous system described by other groups (44, 52) in which the proportion of plasmid-free cells was around 50%.] The parental strain also harbored Δ*tus* and *rpoB*35* mutations, which are expected to enable replication initiated from sites other than *oriC* to progress completely around the chromosome (34).

A similar blue-white screening approach, with *recA*^*+*^ shelter plasmid pHYD5701, was employed to determine viability of Δ*recA* derivatives.

### Genome-wide DNA copy number determinations

Copy number determinations of different genomic regions were performed by whole-genome sequencing (WGS) on phenol-chloroform extracted DNA preparations, essentially as described (36, 53). WGS read data of 200-fold or greater coverage were generated with a paired-end sequencing strategy on an Illumina HiSeq platform.

Sequence reads were aligned to the MG1655 reference genome (NC_000913.3) with the aid of Bowtie2 package (54), following which the *bedcov* function of SAMtools software (55) was used to determine base read counts for successive non-overlapping 1 kb genomic intervals. Base read counts were normalized to the per-kb average of either (i) the aggregate of aligned base read counts, in the case of cultures not subjected to I-SceI cleavage, or (ii) read counts of the 500-kb region between genomic coordinates 683 kb and 1183 kb, for cultures subjected to I-SceI cleavage (since the latter region was expected to be the least affected by DNA resection following such cleavage in the different strains). A second normalization for all cultures was then done against interval-specific normalized read counts of the wild-type strain grown to stationary phase (data for which are shown in Fig. S5A). The moving average method for data smoothening was as described (36), for which the window size and step size used were, respectively, 10 kb and 4 kb. All strains that were used for WGS carried an Δ(*argF-lac*)*U169* deletion, that corresponds to kb coordinates 275 to 373 in the MG1655 reference genome.

The WGS data were also analysed, with the aid of BCFtools (55) and Integrative Genomics Viewer software (56), to validate the different strain genotypes and to exclude presence of significant sequence variations in the cultures.

### Other methods

β-Galactosidase specific activities were determined for *sulA-lac* fusion strains, in cultures grown to mid-exponential phase, by the method of Miller\ (50) and the values are reported in the units defined therein; mean and standard error values were obtained from at least three experiments. The methods for phage P1 transduction (57); recombineering and Flp-recombinase-mediated site-specific excision between FRT sites (48); flow cytometry for SDR detection (58, 59); and chromosome linearization with the aid of N15 phage and engineered *tos* sites (32, 34, 60) were as described. Protocols of Sambrook and Russell (61) were followed for recombinant DNA manipulations, PCR, and transformation.

## Results

### Approaches used to investigate SDR

Three approaches were used in this work to study occurrence and features of SDR in different strains.

(i)As described above, a shelter plasmid-based ‘blue-white screening’ strategy was used initially to identify conditions that serve to rescue Δ*dnaA* lethality (35, 36).

(ii)In a second approach based on flow cytometry, the extent of continued DNA synthesis following addition of a protein synthesis inhibitor spectinomycin was investigated (36, 59). Since DnaA needs to be synthesized continuously for each round of replication initiation at *oriC*, this method provides a quantitation of SDR occurring in individual cells.

(iii)Finally, a WGS approach was used to determine relative copy numbers of different chromosomal regions at the population level in cultures under SDR conditions. Gradients of copy number distribution permit inferences of likely aberrant replication initiation sites (32–34, 36, 37, 58, 62). The WGS approach also serves as a quality check on strain genotypes and possible existence of suppressor mutations.

### Rescue of Δ*dnaA* lethality in a Δ*recD* mutant by imposition of two or more site-specific two-ended DSBs

DNA damage with its accompanying SOS response leads also to a type of SDR that is called inducible SDR [iSDR (4, 63)]. We asked whether iSDR could be co-opted to rescue Δ*dnaA* lethality in particular situations; towards this end, we tested the combined effects of a Δ*recD* mutation and perturbations conferring iSDR.

The Δ*recD* mutation was chosen as candidate for the following reasons. Leach and coworkers had shown excessive DNA synthesis in the vicinity of a site-specific chromosomal DSB in a Δ*recD* mutant (64). Furthermore, copy number determinations by WGS in Δ*recD* strains (62, 64) have revealed a characteristic “mid-terminus peak” that is most commonly associated with mutants exhibiting SDR (5, 32–37, 58, 62, 65).

Viability in absence of DnaA was assessed by blue-white screening assays in strains (in a Δ*tus rpoB*35* background) in which iSDR was provoked after engineering up to four I-SceI cleavage sites in the genome (I-SceI locations are schematically depicted and numbered with Roman numerals in Fig. 1A; primer sequences for recombineering are in Table S2). The derivatives also carried a chromosomal construct for Ara-induced expression of I-SceI enzyme (66). Previous studies have shown that <10% of wild-type cells that have suffered three discrete I-SceI cleavages on the chromosome retain their viability (67), which was also confirmed in our work (see Fig. 1C).

**Figure 1:**
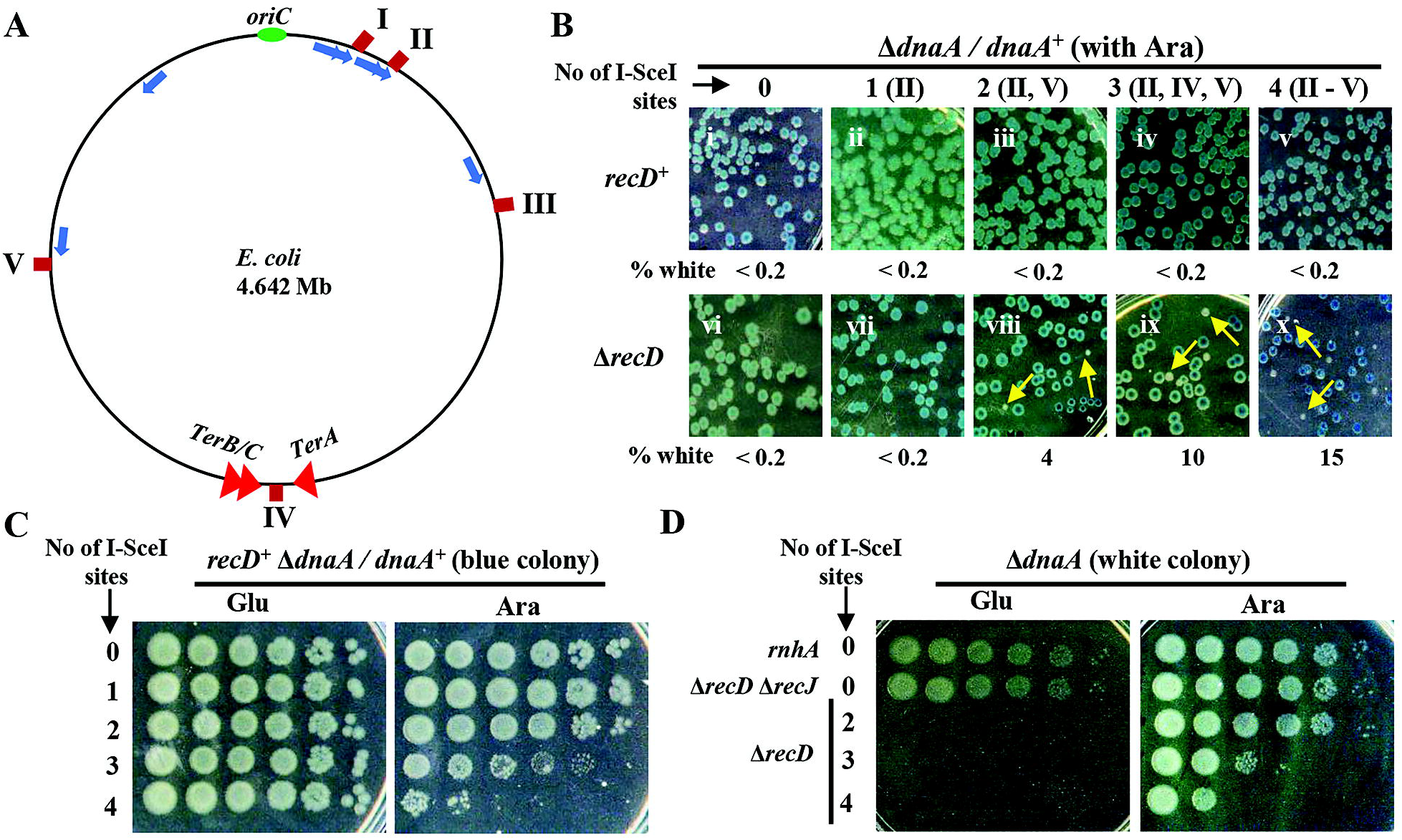
Rescue of Δ*dnaA* lethality by I-SceI cleavages in Δ*recD* mutant. **(A)** Schematic depiction of locations on *E. coli* chromosome of different I-SceI sites used in the study (maroon, marked I to V); also shown are positions of *oriC*, terminus region flanked by *TerA* and *TerB/C*, and the seven *rrn* operons (blue arrows). **(B)** Blue-white assays for *recD*^*+*^ and Δ*recD* derivatives each carrying the indicated number of I-SceI sites (positions denoted in parentheses); all strains (i) were Δ*dnaA* with *dnaA*^*+*^ shelter plasmid pHYD2388, (ii) carried the gene encoding I-SceI enzyme under control of P_*ara*_, and (iii) were plated on Ara-supplemented medium. Representative images are shown, and the numbers beneath each of the panels indicate the percentage of white colonies to the total, that is, viable even in absence of *dnaA*^*+*^ shelter plasmid (minimum of 500 colonies counted). Examples of white colonies are marked by yellow arrows. Strains employed for the different sub-panels were pHYD2388 derivatives of (all strain numbers are prefixed with GJ): i, 16079; ii, 15853; iii, 15857; iv, 15861; v, 15864; vi, 16080; vii, 15888; viii, 15892; ix, 15896; and x, 15899. **(C, D)** Dilution-spotting assays on plates supplemented with 0.2% glucose (Glu) or Ara, of blue or white colonies sourced from plates of panel B; for the top two rows on the plates of panel D, the white colonies were sourced from plates shown, respectively, in Fig. 8 iii and Fig. 4 iii.

The results from these experiments, performed on Ara-supplemented media, indicate that Δ*dnaA* lethality is not rescued (i) even by four sites of I-SceI cleavage in a *recD*^*+*^ strain (Fig. 1B ii-v), nor (ii) by a Δ*recD* mutation by itself or in combination with a single site of I-SceI cleavage (Fig. 1B vi-vii, respectively); Dimude et al (68) have also previously shown that a *recD* mutation fails to suppress lethality in a strain lacking DnaA function. On the other hand, rescue of Δ*dnaA* lethality did occur in Δ*recD* mutants bearing two, three, or four I-SceI cleavage sites (Fig. 1B viii-x). Sustained viability of these Δ*dnaA* derivatives (white colonies) upon sub-culturing was contingent on supplementation of the growth medium with Ara (Fig. 1D), although the proportion of viable cells in strains with three or four cleavage sites was low; a viable Δ*dnaA rnhA* (RNase HI-deficient) derivative (see Fig. 8 iii for a representative blue-white panel) was used as control on these plates (Fig. 1D, top row).

### Two-ended DSBs generated by radiomimetic agents also rescue Δ*dnaA* lethality in Δ*recD* mutant

Radiomimetic agents such as Phleo and Bleo mediate scission of both DNA strands to generate two-ended DSBs (22, 23). When these agents were tested at sublethal doses in the blue-white screening assay (in *recD*^*+*^ or Δ*recD* strains that were Δ*tus rpoB*35*), Δ*dnaA* viability (that is, recovery of white colonies) was noted only with the combination of a Δ*recD* mutation and supplementation with Phleo or Bleo (Fig. 2A, compare v-vi with i-iv). Rescue of Δ*recD* Δ*dnaA* lethality on Phleo-supplemented medium was lost upon introduction of *recD*^*+*^ on a plasmid, but not of the plasmid vector (Fig. 2B). Rescue was also dependent on presence of both Δ*tus* and *rpoB*35* mutations in the strain (Fig. S1A). Furthermore, as expected, the Δ*recD* white colonies obtained on Phleo-supplemented medium retained viability only on medium supplemented with sublethal Phleo (Fig. 2C, bottom row; Δ*dnaA rnhA* colony was used as control, top row).

**Figure 2:**
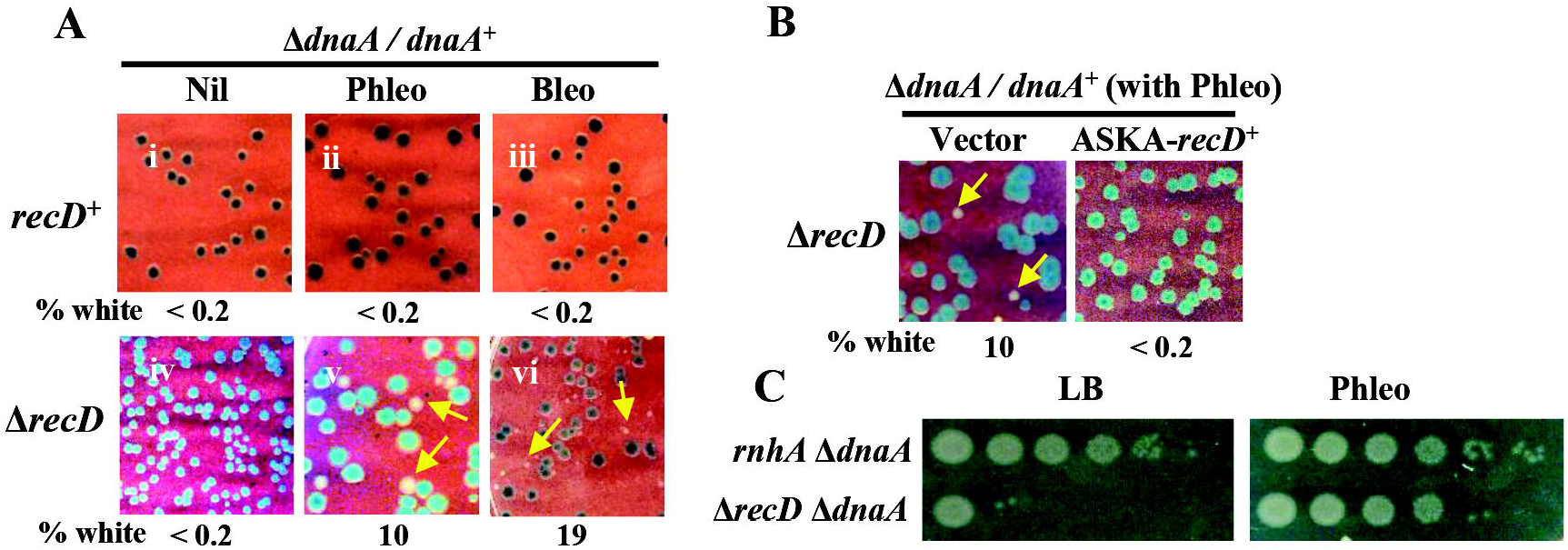
Rescue of Δ*dnaA* lethality by Δ*recD* in presence of sublethal Phleo or Bleo. **(A, B)** Blue-white assay results with pHYD2388 derivatives of GJ18607 (*recD*^*+*^) or GJ15935 (Δ*recD*) are depicted as described in legend to Fig. 1B, on plates without or with Phleo or Bleo supplementation. In panel B, the Δ*recD* strain also carried the ASKA plasmid vector (pCA24N) or the latter’s *recD*^*+*^ derivative. **(C)** Dilution-spotting assays on medium without or with Phleo supplementation, of white colonies from plates shown in Fig. 8 iii (for top row) and Fig. 2A v (for bottom row).

Accordingly, we conclude that a discrete number of two-ended DSBs, generated by either radiomimetic agents or site-specific endonucleolytic cleavages, are necessary and sufficient to rescue Δ*dnaA* lethality in a Δ*recD* mutant.

### Perturbations that generate one-ended DSBs do not rescue Δ*dnaA* lethality in Δ*recD* strain

Perturbations that generate nicks or abasic sites in DNA lead subsequently to collapse or disintegration of replication forks and thereby to one-ended DSBs on the chromosome. Such perturbations include mutations in genes *rdgB, polA*, or *dut* (12, 69, 70), or exposure to genotoxic agents such as norfloxacin, ciprofloxacin, or mitomycin C (10, 12, 71). One- and two-ended DSBs are similar to one another in that (i) both provoke the SOS response and iSDR (4, 63, 72–74), and (ii) their repair is RecA- and RecBCD-dependent (9, 12, 14, 19, 74).

When one-ended DSB-generating perturbations were tested in a Δ*recD* mutant (carrying Δ*tus* and *rpoB*35* alleles), none was able to confer Δ*dnaA* lethality rescue (Fig. 3A). For each of the exogenous genotoxic agents used, we showed that there is >10^5^-fold killing of cells of a Δ*recA* derivative at the same concentration (Fig. S2A; Phleo supplementation was used as positive control in this experiment), implying that DSBs are indeed being generated in all cells under these conditions (12). As expected, the Δ*recD* derivative was as tolerant as the wild-type parent to all the agents (Fig. S2A). Measurements of β-galactosidase specific activity, in derivatives bearing an SOS-responsive *sulA-lac* fusion (75), indicated that the magnitude of SOS induction by one-ended and two-ended DSB damage was similar (around 4-fold), and that this was true for both wild-type and Δ*recD* strains (Fig. 3B).

**Figure 3:**
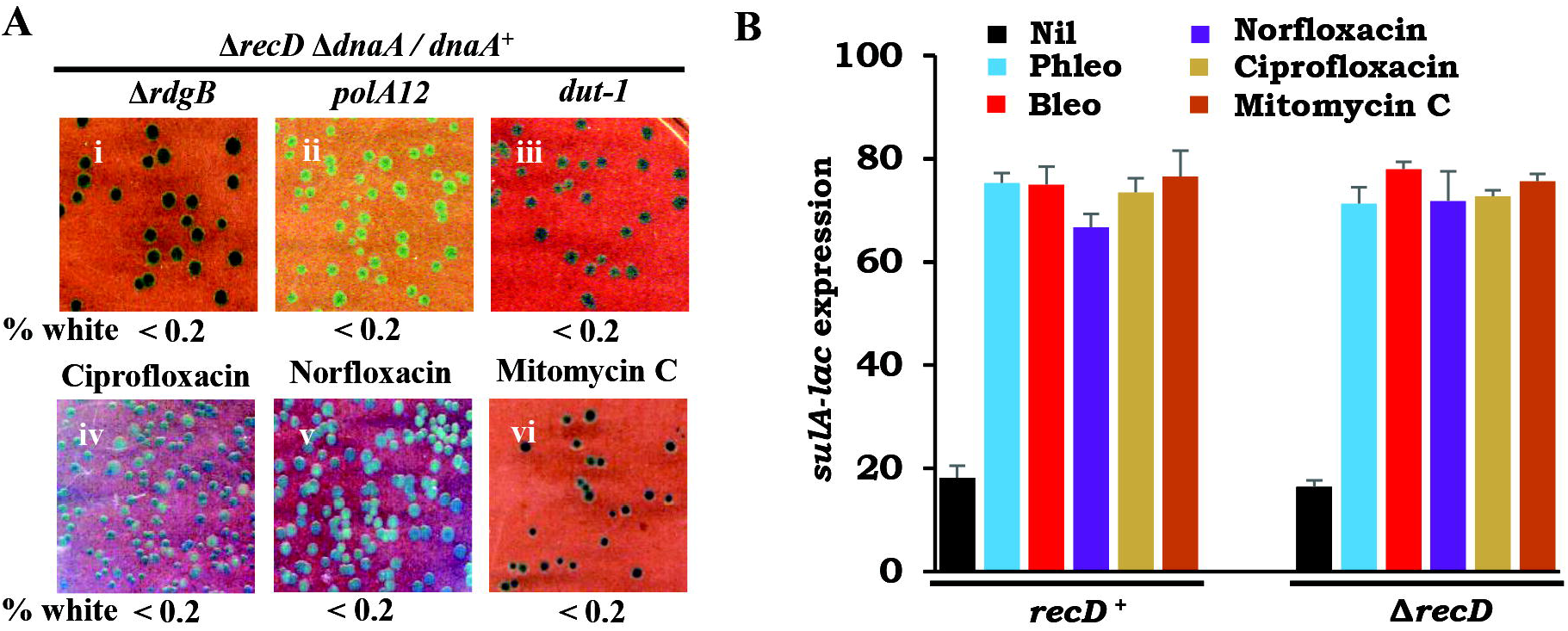
Δ*dnaA* lethality is not rescued in Δ*recD* strain by perturbations generating one-ended DSBs. **(A)** Blue-white assay results are depicted, as described in legend to Fig. 1B, for Δ*recD* derivatives subjected to one-ended DSBs through either additional mutations (panels i-iii) or exposure to genotoxic agents (panels iv-vi), as indicated. Concentrations used were (μg/ml): ciprofloxacin, 0.005; norfloxacin, 0.02; and mitomycin C, 1. Strains used for the different panels were pHYD2388 derivatives of the following (all strain numbers are prefixed with GJ): i, 15939; ii, 15940; iii, 16082; and iv-vi, 15936. **(B)** β-Galactosidase specific activity (Miller units ± SE) in *sulA-lac* strains GJ16091 (*recD*^*+*^) and GJ16092 (Δ*recD*) without and with exposure to agents causing one-ended DSBs (norfloxacin, ciprofloxacin or mitomycin C at concentrations indicated above) or two-ended DSBs (Phleo or Bleo).

The results therefore indicate that one- and two-ended DSBs behave dissimilarly in the SDR assay with Δ*recD* strains. Furthermore, they suggest that iSDR is necessary but not by itself sufficient for rescue of Δ*dnaA* lethality in Δ*recD* mutants, since neither one-ended DSBs, nor a two-ended DSB generated at a single I-SceI cleavage site, confer viability to a Δ*recD* Δ*dnaA* derivative although they too elicit iSDR (4, 63, 72, 73).

### Rescue of Δ*dnaA* lethality by a combination of Δ*recD* and Δ*recJ* mutations

We then asked whether any other single gene mutation, which by itself cannot rescue Δ*dnaA* lethality, can do so in combination with Δ*recD*. Of a variety of candidate genes tested, a Δ*recJ* mutation (associated with deficiency of RecJ 5’ ss-DNA exonuclease) was shown to synergize with Δ*recD* to confer viability to a Δ*dnaA* mutant (Fig. 4A, compare iii with i-ii; see also Fig. 1D, row 2); the phenotype was complementable by plasmids expressing RecD or RecJ, but not by the empty vector (Fig. 4B, compare ii-iii with i). On the other hand, the combination of Δ*recD* with loss of any one of the following DNA nucleases Exo I, Exo VII, or SbcCD (encoded by *xonA, xseA* and *sbcCD*, respectively) could not rescue Δ*dnaA* lethality (Fig. S1B).

**Figure 4:**
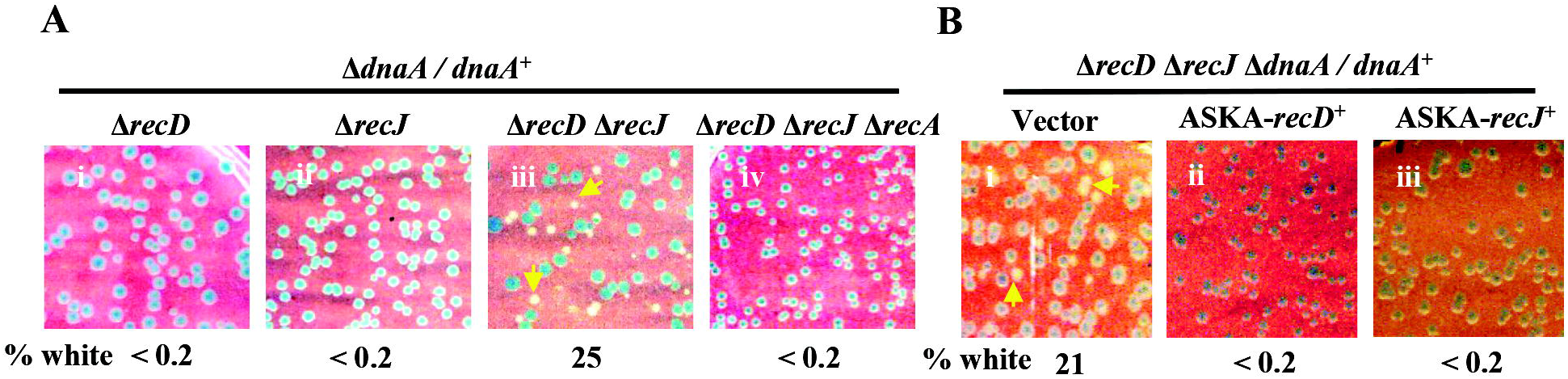
Rescue of Δ*dnaA* lethality by Δ*recD* Δ*recJ*. **(A, B)** Blue-white assay results are depicted, as described in legend to Fig. 1B, for derivatives whose relevant genotypes are given on top (all were Δ*dnaA/dnaA*^*+*^). Strains used for different sub-panels of panel A were pHYD2388 derivatives of the following (all strain numbers are prefixed with GJ): i, 15936; ii, 16083; iii, 15949; and iv, 16084. In panel B, the pHYD2388 derivative of Δ*recD* Δ*recJ* strain GJ15949 also carried the ASKA plasmid vector (pCA24N) or the latter’s *recD*^*+*^ or *recJ*^*+*^ derivative.

Suppression of Δ*dnaA* inviability, both in the Δ*recD* Δ*recJ* strain and in Δ*recD* with Phleo, was abolished upon introduction of a *priA300* mutation that leads to loss of helicase activity of PriA (Fig. S3, i-ii); PriA helicase may regulate abnormal replication restart in certain mutant backgrounds (34, 52, 76), but it is also postulated to be required for one or more of the redundant restart pathways in a wild-type strain (9, 19, 20) and its loss is correlated with compromised SDR (77). Another candidate mutation that was tested Δ*dinG* also abolished viability of Δ*dnaA* both in the Δ*recD* Δ*recJ* strain and in Δ*recD* on Phleo-supplemented medium (Fig. S3, iii-iv). The DinG helicase is required to facilitate progression of replisomes when they are in head-on conflict with *rrn* operon transcription (such conflicts are expected in SDR), presumably by catalyzing removal of RNA-DNA hybrids generated at the sites of conflict (24, 78); we could show that RNase HI overexpression (achieved from an ectopic chromosomal P_*ara*_*-rnhA* construct) restores Δ*dnaA* rescue in the Δ*dinG* derivatives above (Fig. S3, vii-viii), whereas the control construct lacking *rnhA* was unable to do so (Fig. S3, v-vi).

### No evidence for spontaneous DSBs in *recD recJ* mutant

Our findings that Δ*dnaA* lethality rescue is achieved with multiple two-ended DSBs in Δ*recD*, and separately so by combined loss of both RecD and RecJ, raised the question whether occurrence of spontaneous two-ended DSBs is responsible for the phenotype in the latter. A number of studies have shown that viability of a *recD recJ* mutant is more or less similar to that of the wild-type parent in absence of exposure to DNA damaging agents; however, it is moderately defective for homologous recombination during conjugation or transduction, and it is very sensitive to DNA damage by various exogenous agents as also with the *dut-1* mutation (79–85).

By employing a *recA*^*+*^ shelter plasmid in the blue-white screening assay, we could show that loss of RecA does not confer lethality or even any greater sickness in the Δ*recD* Δ*recJ* derivative than in the wild-type or cognate single mutant strains [Fig. S2B, compare relative sizes of blue (*recA*^*+*^) and white (Δ*recA*) colonies in different panels]. This serves to exclude occurrence of spontaneous DSBs in the Δ*recD* Δ*recJ* mutant, since a *recA* mutant cell is killed with even one DSB on the chromosome (12, 13, 42, 53, 86). The triple mutant *recD recJ recA* of *Salmonella typhimurium* is also viable (84). It would appear, therefore, that the combination of *recD* and *recJ* mutations does not by itself cause damage to DNA, although it does render cells deficient for repair of DNA lesions generated by genotoxic perturbations.

On the other hand, rescue of Δ*dnaA* lethality was not observed in the triple mutant Δ*recD* Δ*recJ* Δ*recA* (Fig. 4 iv), indicating that SDR in the Δ*recD* Δ*recJ* strain is RecA-dependent.

### Demonstration by flow cytometry of SDR in Δ*recD* derivatives

In the flow cytometric method for SDR detection, incorporation of the nucleotide analog EdU (as detected with Alexa Fluor 488 staining) is used as a measure of DNA synthesis in individual cells after protein synthesis is inhibited by addition of spectinomycin (36, 59). The experiments were performed in Δ*tus rpoB*35* derivatives of different strains; we did not, however, employ the method for cultures that had been subjected to overt DNA damage (such as with Phleo or Bleo exposure, or I-SceI cleavage), since such damage is known to trigger iSDR (4).

EdU incorporation in absence of spectinomycin was uniformly high for all the strains tested (Fig. S4). Following spectinomycin treatment, the proportion of cells exhibiting DNA synthesis above threshold was (in percent): wild-type (negative control), 6; *rnhA* (positive control), 88; Δ*recJ*, 29; Δ*recD*, 36; Δ*recD* Δ*recJ*, 85; and Δ*recD* Δ*recJ* Δ*recA*, 25 (Fig. 5 i-vi, respectively). Of these, as described above, it is only the Δ*recD* Δ*recJ* derivative that is viable in absence of DnaA, and its EdU positivity was comparable to that in the positive control.

**Figure 5:**
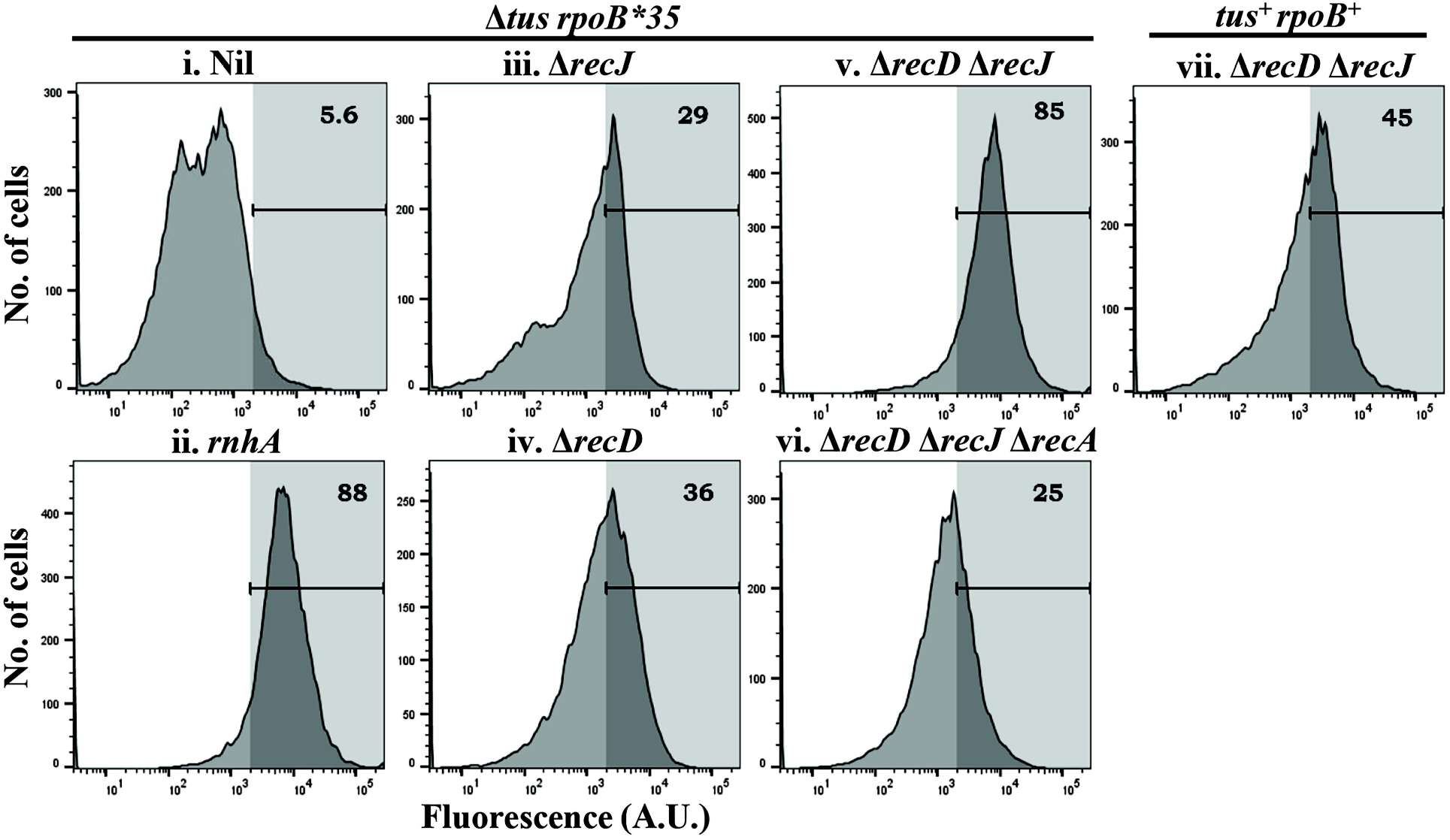
Demonstration by flow cytometry of SDR in Δ*recD* Δ*recJ* mutants. DNA synthesis was measured as described (29, 45), for different strains (genotype given on top of each panel) after addition of spectinomycin to the cultures. In each panel, the percentage of cells in which the fluorescence value exceeded the threshold (denoted by vertical line) set at 2 × 10^3^ arbitrary units (A.U.) is given at top right. Strains used for the different panels were (all strain numbers are prefixed with GJ): i, 18609; ii, 15981; iii, 15985; iv, 15984; v, 15986; vi, 15987; and vii, 15979.

For the Δ*recD* Δ*recJ* mutant, we also determined the proportion of EdU-positive cells after spectinomycin treatment of the *tus*^*+*^ *rpoB*^*+*^ derivative, in order to assess the quantitative contribution to SDR of the mutations in these genes (Fig. 5 vii). At 45%, this proportion was approximately one-half of that in the isogenic Δ*tus rpoB*35* strain.

Taken together, these data indicate that (i) substantial SDR occurs even in a Δ*recD* mutant without overt DNA damage; (ii) in a Δ*recD* Δ*recJ* mutant, loss of RecA is associated with reduction of biochemical SDR, which is consistent with the genetic assay results that the triple mutant does not support Δ*dnaA* viability; and (iii) presence of Δ*tus* and *rpoB*35* mutations is correlated with increased EdU incorporation in cells inhibited for protein synthesis. The last observation serves to re-affirm the proposal (34) that replication initiated from sites other than *oriC* is impeded for its progression around the chromosome by Tus-bound *Ter* sites and by conflicts between transcription and replication. Our data also support the notion advanced earlier (4) that the genetic assay imposes a far more stringent threshold for detection of SDR than do the biochemical assays.

### Genome-wide DNA copy number determinations in Δ*recD* derivatives with Phleo supplementation or with Δ*recJ* mutation

As mentioned above, replication initiation from *oriC* in asynchronously growing cultures is characterized by a bidirectional *oriC*-to-*Ter* gradient of copy number distributions, with this pattern being disturbed in mutants exhibiting SDR (5). We performed WGS experiments in different Δ*recD* or Δ*recD* Δ*recJ* derivatives to determine, at the population level, the nature of such alterations and thus to obtain insights into SDR mechanisms operating in them.

Given that rescue of Δ*dnaA* lethality, as a manifestation of SDR, had been observed in this study for the Δ*recD* mutant with Phleo and for the double mutant Δ*recD* Δ*recJ*, the WGS experiments were performed for both these situations, using the *dnaA*^*+*^ *tus*^*+*^ *rpoB*^*+*^ derivatives; the control cultures were the wild-type strain as well as its single mutant Δ*recJ* and Δ*recD* derivatives (all *dnaA*^*+*^ *tus*^*+*^ *rpoB*^*+*^, without Phleo). Copy number distribution by WGS were also determined for the cultures that were viable in absence of DnaA (that is, Δ*dnaA* Δ*tus rpoB*35* Δ*recD* with Phleo, and Δ*dnaA* Δ*tus rpoB*35* Δ*recD* Δ*recJ*) as well as for their cognate *dnaA*^*+*^ derivatives.

The cultures for wild-type as well as its Δ*recD* and Δ*recJ* single mutant derivatives exhibited the expected bidirectional *oriC-*to-*Ter* gradient in their DNA copy number distributions (Fig. 6 i-ii, and Fig. S5B, respectively). As mentioned above, one copy number feature that is common across several perturbations which serve to restore viability in absence of DnaA function, is presence in the cognate *dnaA*^*+*^ cultures of a “mid-terminus peak” [that is, an increased copy number of DNA of the terminus region in *dnaA*^*+*^ *tus*^*+*^ *rpoB*^*+*^ derivatives (32–37, 58, 62, 65)]. We have previously suggested that the mid-terminus peak represents a population averaging of copy number contributions from superimposed aberrantly initiated replication forks that have then been trapped between the Tus-bound *Ter* sites of the terminus (5). We note that this feature was preserved also for the two sets of perturbations (Δ*recD* with Phleo, and Δ*recD* Δ*recJ*) that were correlated with SDR in the present study (Fig. 6 iii-iv, respectively). The peak was not observed in the isogenic Δ*tus rpoB*35* derivatives (Fig. 6 v-vi, respectively), which is consistent with the proposal (33, 34) that it arises because of fork-trapping at *Ter* sites to which Tus is bound; the copy number gradient in these derivatives was also flattened which, as discussed below, supports the notion that SDR in these cells is sufficiently robust to counteract the *oriC*-directed copy number gradient averaged from the population.

**Figure 6:**
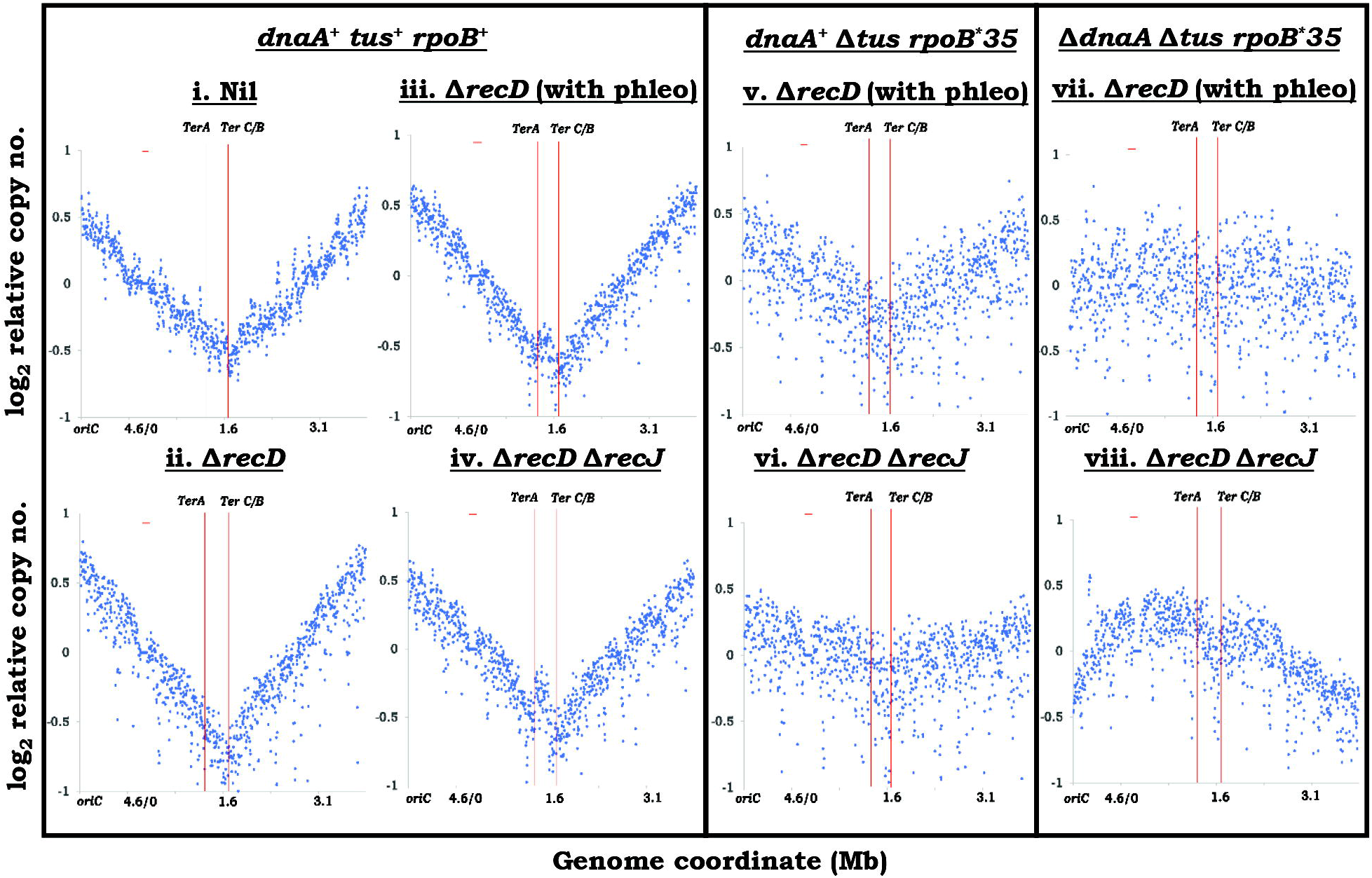
Chromosomal DNA copy number analysis by WGS. Distinguishing strain genotypes and Phleo supplementation were as indicated on top of the panels. DNA copy numbers are plotted (after normalization) as moving averages across the genome. For these graphical representations, the circular 4642-kb long chromosome is shown linearized at *oriC*, with genome coordinates on the abscissa corresponding to the MG1655 reference sequence (wherein *oriC* is at 3926 kb); the positions of *TerA* and *TerC/B* are marked. Strains used for the different panels were (all strain numbers are prefixed with GJ): i, 18601, ii-iii, 15988; iv, 15991; v, 15984; vi, 15986; vii, 15936; and viii, 15949.

For the pair of Δ*dnaA* cultures (which were also Δ*tus rpoB*35*), absence of an *oriC* peak was as expected; the copy number distribution was more or less flat across the genome in the Δ*recD* derivative cultured with sublethal Phleo (Fig. 6 vii), whereas that in the Δ*recD* Δ*recJ* mutant exhibited a broad peak in the vicinity of the terminus region (Fig. 6 viii). In the Discussion below, these patterns are interpreted in the context of different sites of postulated aberrant replication initiation in the cultures.

### Genome-wide DNA copy number determinations in Δ*recD* derivatives with site-specific two-ended DSBs

Since we had identified that Δ*dnaA* lethality can be rescued in a Δ*recD* strain also by imposition of two-ended DSBs at two or more discrete sites through I-SceI cleavage (in presence of Δ*tus* and *rpoB*35* mutations), we determined by WGS the DNA copy number distributions in four of these Δ*dnaA*-suppressed derivatives: three bearing different combinations of dual I-SceI sites (of which site II near *dusA* was common and the second numbered site I, III, or V was near *yihQ, yaiZ*, and *nadB*, respectively), and one with three I-SceI sites (II, IV and V; site IV being near *yddT*). All cultures were grown in continuous presence of Ara since this was a requirement for their viability.

As controls, *dnaA*^*+*^ derivatives of the *recD*^*+*^ and Δ*recD* strains each carrying two I-SceI cleavage sites (II and V) were used in the WGS experiments after growth in continuous presence of Ara (both were also Δ*tus* and *rpoB*35*). Previous studies have shown that sustained cleavage in the wild-type strain at a single chromosomal I-SceI site results in attainment of an equilibrium between DNA resection on the one hand and DNA repair on the other, such that an asymmetric V-shaped pattern of reduction in DNA copy number is generated around the cleavage site; this dip in copy number extends to approximately 100 kb towards *oriC* and 200 kb towards *Ter* (53, 86).

The results from these population-level DNA copy number determinations are shown in Figure 7. In the *dnaA*^*+*^ *recD*^*+*^ control strain (Fig. 7 i), copy number reductions were observed in the vicinity of both I-SceI cleavage sites II and V, but its magnitude was apparently less than that described in the previous studies for a single site of cleavage (53, 86); it is possible that either cleavage by I-SceI, or resection by RecBCD, becomes rate-limiting with more than one I-SceI site on the chromosome. In the *dnaA*^*+*^ Δ*recD* derivative (Fig. 7 ii), copy number reductions at the I-SceI sites (II and V) were less than those in the *recD*^*+*^ control, which is similar to that also described earlier for single-site cleavage (86).

**Figure 7:**
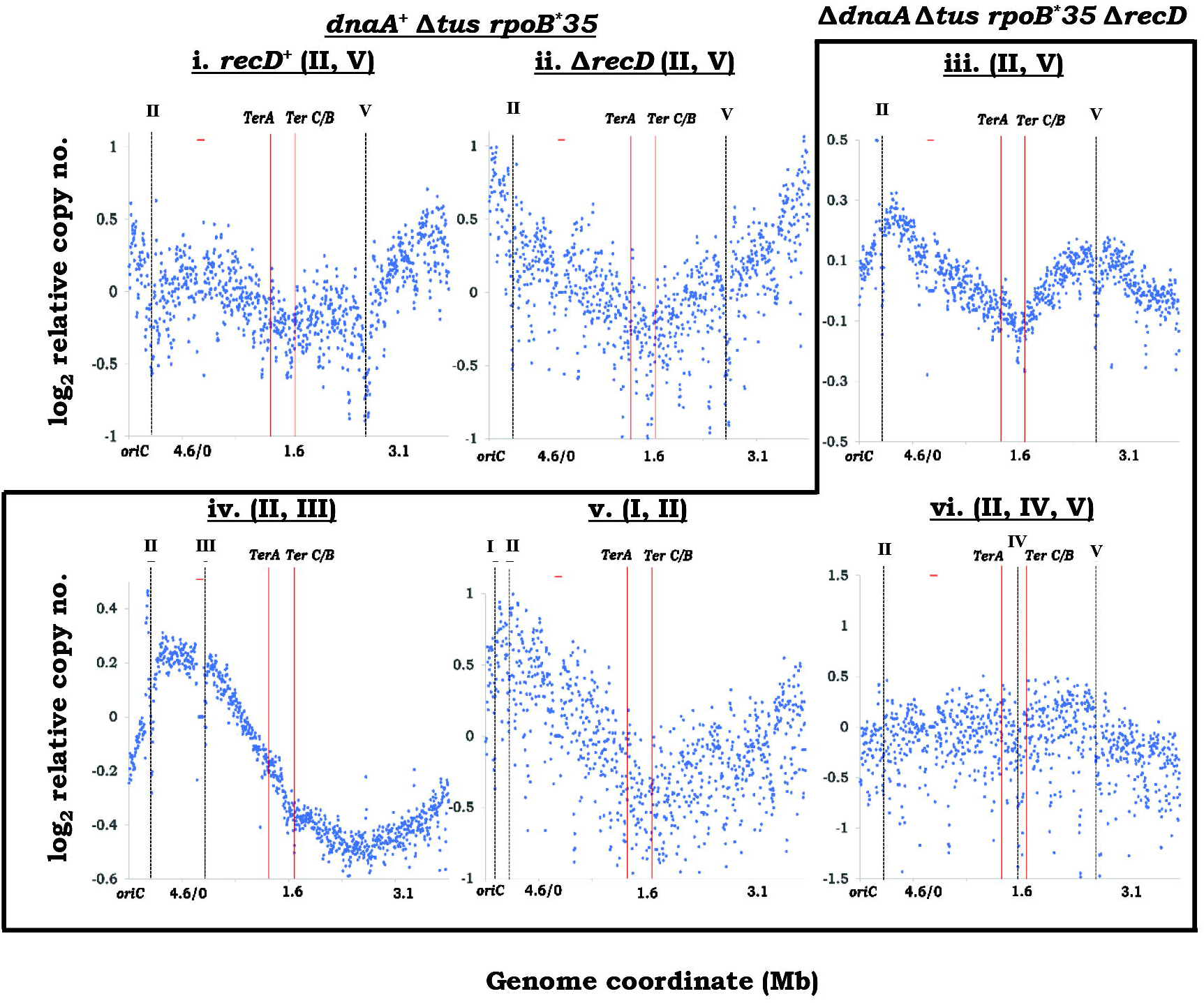
Chromosomal copy number distributions, as determined by WGS, in Δ*dnaA* Δ*recD* strains rendered viable by I-SceI cleavages. Distinguishing strain genotypes are indicated on top of the panels, and Roman numerals within parentheses denote the distinct I-SceI sites as shown in Fig. 1A; all strains also carried the construct for Ara-inducible expression of I-SceI enzyme, and were cultured in continuous presence of Ara for these experiments. Representations of WGS analysis data and notations used are as in the legend to Fig. 6; positions of I-SceI sites are also marked in each of the panels. Strains used for the different panels were (all strain numbers are prefixed with GJ): i, 15857/pHYD2388; ii, 15892/pHYD2388; iii, 15892; iv, 15889; v, 15890; and vi, 15896.

In the Δ*dnaA* Δ*recD* cultures rendered viable by I-SceI cleavages at various sites, DNA copy number patterns were consistent with the idea that aberrant replication was being initiated in the vicinity of the cleavage sites. Thus, in a derivative with the dual cleavage sites II and V (Fig. 7 iii), two distinct copy number peaks were observed, whereas in that with sites II and III (Fig. 7 iv), a broad peak was located approximately midway between the two loci; and in a derivative with sites I and II (which are very close to one another, as also to *oriC*), the copy number gradient resembled that for *oriC-* initiated replication itself (Fig. 7 v). Finally, in a Δ*dnaA* strain with three I-SceI sites (II, IV, V) that were somewhat evenly separated on the chromosome, the copy number distribution was approximately flat across the genome (Fig. 7 vi), which is reminiscent of the pattern previously reported in derivatives of the archaeon *Haloferax volcanii* in which all replication origins were deleted, and chromosome duplication was achieved through DNA synthesis triggered by homologous recombination (87). The interpretations from these data with respect to the mechanism of SDR in absence of RecD are provided in the Discussion.

### Effect of chromosome linearization on rescue of Δ*dnaA* lethality in Δ*recD* mutants

*E. coli* cells whose chromosome has been linearized in its terminus region by use of the phage N15-*tos* system retain viability without any obvious fitness deficit (60). In the context of SDR, it has been suggested (based on studies with N15-*tos*) that when the source of aberrant replication initiation is itself in the terminus region (as has been postulated for *recG* or 3’-exonuclease mutants), viability in absence of DnaA function is exhibited only if the chromosome is circular (32, 34, 37); on the other hand, RNase HI deficiency confers viability even in linear chromosome derivatives lacking DnaA (32).

With the aid of the blue-white screening assay, the effect of chromosome linearization by N15-*tos* on rescue of Δ*dnaA* lethality in different derivatives was tested. Our results support earlier findings (32, 34) that Δ*dnaA* lethality rescue (in a Δ*tus rpoB*35* background) by Δ*rnhA* is retained but that by Δ*recG* is abolished after chromosome linearization (Fig. 8, compare, respectively, iii with iv and v with vi); they also show that with the linear chromosome constructs, Δ*recD* with Phleo, but not Δ*recD* Δ*recJ*, is able to confer Δ*dnaA* viability (Fig. 8, compare, respectively, vii with viii and ix with x).

**Figure 8:**
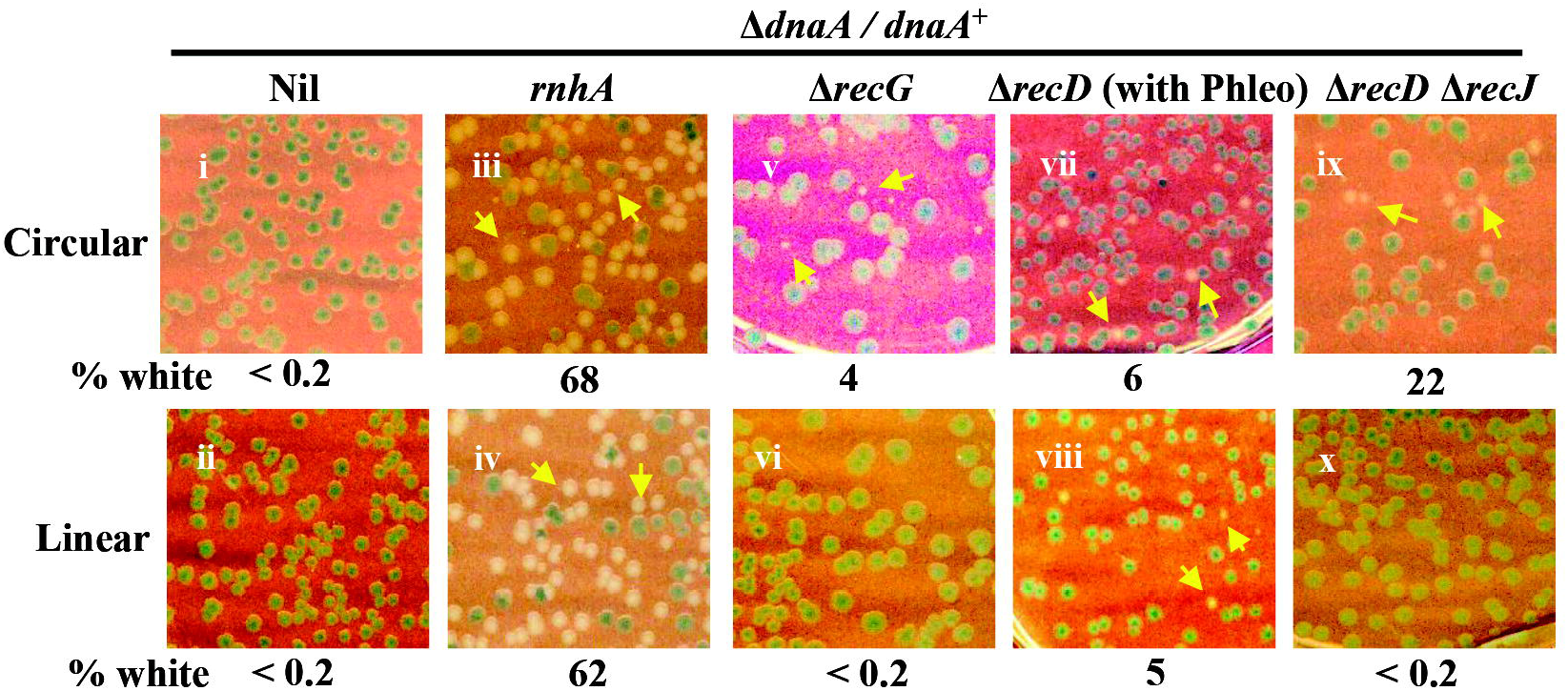
Effect of chromosome linearization on rescue of Δ*dnaA* lethality by different perturbations. Blue-white assay results are depicted, as described in legend to Fig. 1B, for isogenic pairs of circular- and linear-chromosome strains (upper and lower rows, respectively) with relevant perturbations as indicated on top. The proportion of white colonies to total was consistently elevated in *rnhA* mutants compared to that in other strains (see also Fig. 9), the reasons for which remain to be determined. Strains used for the different panels were pHYD2388 derivatives of the following (all strain numbers are prefixed with GJ): i, 18607; ii, 15993; iii, 16081; iv, 15995; v, 16464; vi, 15999; vii, 15936; viii, 16001; ix, 15949; and x, 16003.

## Discussion

The roles of the dual function RecBCD helicase-nuclease for recombinational repair of one-ended and two-ended DSBs have been well characterized; the enzyme catalyzes resection of the DSB end and then mediates formation of a RecA nucleoprotein filament on ss-DNA (12–17). RecBCD’s actions have also been implicated in the handling of reversed replication forks (88, 89) as well as in fidelity of chromosomal terminus region inheritance (90–93).

In comparison to the RecBCD enzyme, the RecBC complex (without RecD subunit) is attenuated for nuclease activity, even as it retains helicase activity and is proficient for DSB repair (12, 17). Reduced resection activity on ds-DNA ends in a *recD* mutant has recently been confirmed by in vivo imaging experiments (94), and this property underpins several phenotypes of *recD* mutants such as (i) generation of linear plasmid multimers (62, 95, 96), and (ii) ability to integrate linear DNA fragments in chromosomal recombineering (97, 98).

Chromosomal DNA over-replication in *recD* mutants has also been recorded earlier in two situations, namely in the chromosomal terminus region (62, 64) and in the vicinity of a DSB generated at an engineered palindromic sequence on the lagging strand template behind a replication fork (64). For the latter, a model to explain the phenomenon of DNA amplification in the Δ*recD* mutant has been proposed that invokes uncoordinated invasion of the two DSB ends into an intact sister chromosome (64). The present study demonstrates that DNA over-replication in cells attenuated for RecBCD’s nuclease activity can indeed be sufficient to confer viability in absence of *oriC*-dependent replication, as further discussed below.

### Suppression of Δ*dnaA* lethality in Δ*recD* mutants, and its relationship to iSDR

We have shown here that in strains carrying the Δ*tus* and *rpoB*35* mutations, Δ*dnaA* inviability is suppressed by a Δ*recD* mutation in two situations: (i) when two-ended DSBs are generated, either with sublethal Phleo or Bleo or by two or more site-specific endonucleolytic cleavages mediated by I-SceI; or (ii) in absence of another DNA exonuclease RecJ. The former suppression is exhibited in strains with a circular or a linear chromosome, whereas the latter occurs only in strains with a circular chromosome. One-ended DSBs in the Δ*recD* mutant do not support Δ*dnaA* viability.

If the phenotypes above are viewed in context of iSDR, a reasonable interpretation would be that iSDR is responsible for Δ*dnaA*-independent survival following generation of two or more two-ended DSBs in a Δ*recD* mutant. Asai et al (99) have shown that the magnitude of iSDR is elevated in *recD* or *recJ* single mutants following DNA damage.

Nevertheless, iSDR may by itself not be a sufficient explanation for Δ*dnaA* suppression in the Δ*recD* strain, since one-ended DSBs (or a single two-ended DSB) are unable to suppress Δ*dnaA* lethality although they too provoke iSDR (4, 63, 72, 73); our results also indicate that SOS induction in a Δ*recD* mutant following one- or two-ended DSBs is quantitatively similar. Furthermore, for the double mutant Δ*recD* Δ*recJ*, RecA-independence for viability suggests that its DNA has not suffered any damage, and therefore that rescue of Δ*dnaA* lethality in this situation is unrelated to iSDR.

### Model of rolling-circle replication at merger sites of convergent replication forks

The model which we propose to explain the findings above draws on those outlined earlier by the groups of Courcelle (62, 93, 100, 101) and of Rudolph (34, 37, 102). In wild-type *E. coli*, the common feature between chromosomal terminus region replication on the one hand and ends-in replication during two-ended DSB repair on the other is that of convergent replication forks which subsequently merge; such fork mergers are not a feature of one-ended DSB repair. An example of such converging fork configurations at the chromosomal terminus region is depicted in Figure 9A-B, but equivalent structures would also occur at sites of two-ended DSB repair.

**Figure 9:**
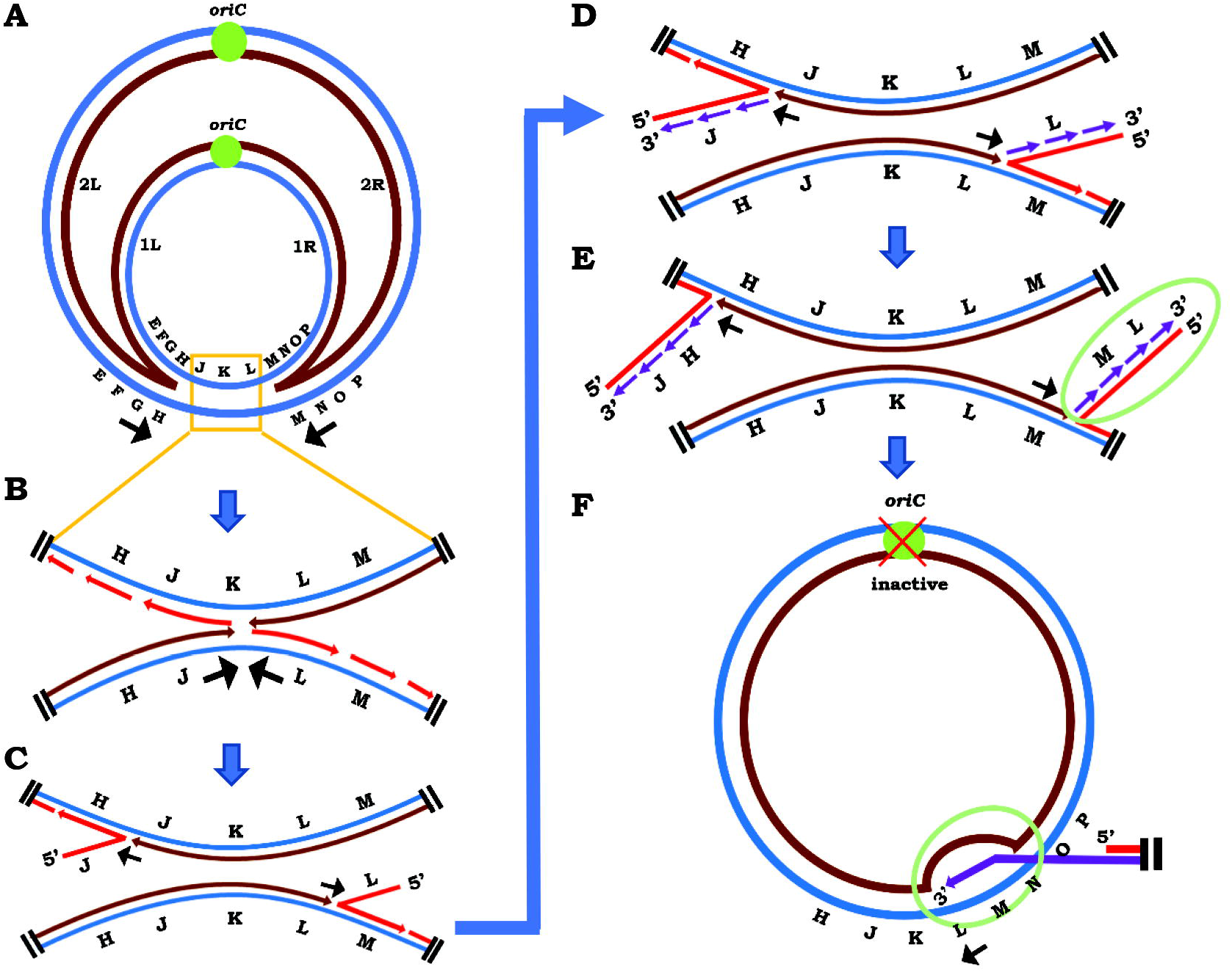
Model to explain SDR in strains lacking RecD and RecJ. **(A)** Schematic depiction of converging replication forks in chromosomal terminus region. Positions and directions of replisome progression (in all panels) are marked by black arrows. The two replichore pairs are designated as 1L-1R and 2L-2R, respectively; and the letters E through P are used to denote different segments of the *oriC-*distal chromosomal region. **(B)** Zoom-in of the replicated terminus region of panel A, wherein disposition and polarity of DNA strands is also shown. **(C)** Progression of components of one replisome past those of the other is postulated to result, ultimately, in generation of a 5’-flap (red) on the opposite lagging strand (shown as a mirror-symmetric pair, although it is possible that the two individual events occur in different sub-populations of cells). **(D, E)** Assembly of replication restart machinery at the fork structures of panel C results in replisome establishment for Okazaki fragment synthesis and initiation of a type of rolling-circle (σ-mode) replication. **(F)** Homologous recombination of the linear extruded duplex (marked by the green oval in panel E) with a sister chromosome to establish a second replisome for *oriC-* and DnaA-independent replication. Please see text for details.

We suggest that such converging replisomes may cross one another, as was postulated earlier by the Rudolph and Courcelle labs (37, 93) and has also been shown to occur in eukaryotic systems (103). We further propose that this ultimately results in a structure in which an extended leading strand has displaced the lagging strand of the opposite fork to create a 5’ flap, as shown in Figure 9C.

Should this model be correct, the intermediate processing steps between the structures depicted in panels B and C of Figure 9 would remain to be determined, as too the roles if any of different DNA helicases (DnaB, RecG, PriA), DNA polymerases (I, III), and DNA topoisomerases (gyrase, topoisomerases III and -IV) in mediating these events. It has been suggested earlier that when replisomes cross one another at a site of fork convergence, 3’ flaps are initially generated which can then be converted to 5’ flaps by action of RecG helicase (34, 37).

According to our model, the 5’ flap that is generated then serves as template for synthesis of Okazaki fragments (Fig. 9D), similar to that occurring during rolling circle replication of λ phage in the late phase of its lytic cycle (104, 105); this would require replication restart mechanisms mediated by PriA/B/C to aid in replisome assembly at the 3-way junction. The linear DNA duplex so generated will ordinarily be rapidly degraded by RecBCD nuclease in a wild-type strain; however, in a Δ*recD* mutant which exhibits attenuated nuclease activity on ds-DNA ends (12, 17, 94, 97, 98), it is expected both to be reasonably stable and to undergo further extension (as shown in Fig. 9E). Thus, the “driver” of *oriC-* and DnaA-independent replication under these conditions is establishment of rolling-circle replication at a site where a pair of converging replication forks merge.

The linear duplex that is generated by rolling-circle replication may also undergo homologous recombination with a circular sister chromosome within the cell, which is especially likely to be favoured in *recD* mutants since the latter are hyper-recombinogenic (85). Such an event would then set up a second “driver” of DnaA-independent chromosomal replication (Fig. 9F), which together with the first “driver” of rolling-circle replication described above will mediate bidirectional replication of chromosomal DNA from a site of fork merger (note locations and opposing directions of replisome progression in panels E and F of Fig. 9). Our findings in the WGS studies, of bidirectional gradient of DNA copy number from I-SceI cleavage sites, may accordingly be explained.

Bidirectionality of replication is also implicit from the mirror-symmetric disposition of the pair of 5’-flap structures that are postulated at sites of fork merger (as shown in Fig. 9C). However, it is quite likely that this component of bidirectionality is merely a population phenomenon, and that it may not be a simultaneous occurrence in individual cells.

Our model would also explain several other observations from this and earlier studies. (i) Since the 5’ flap is susceptible to RecJ-mediated degradation (which can happen prior to synthesis of the first Okazaki fragment), the extrusion process will be expected to be more efficient in a Δ*recD* Δ*recJ* mutant. Consequently, such synthesis from the chromosomal terminus region alone may be sufficient to support Δ*dnaA* viability in this mutant whereas in a Δ*recD recJ*^*+*^ strain, additional converging fork pairs (at sites of two-ended DSB repair) would also be required for the purpose. (ii) As a corollary, since the locus of additional DNA synthesis is restricted to the terminus region of Δ*recD* Δ*recJ* mutants, it is rendered sensitive to chromosome linearization (32, 34, 37); on the other hand, since such synthesis is distributed across several sites in Δ*recD* mutants that have suffered two-ended DSBs, rescue of Δ*dnaA* lethality is observed even in the linear chromosome derivatives. (iii) The same features would also explain, respectively, the terminus region peak and flattened DNA copy number distribution patterns in cultures of Δ*dnaA* Δ*recD* Δ*recJ* and Δ*dnaA* Δ*recD* with Phleo. (iv) Progression of many of the “driver” replisomes would occur in a retrograde *Ter*-to-*oriC* direction, necessitating presence of the Δ*tus* and *rpoB*35* mutations to achieve replication of the entire chromosome in absence of DnaA. (v) Abolition of Δ*dnaA* rescue in Δ*recD* derivatives with *priA300* mutation can be related to compromise by the latter of one or more replication restart pathways (9, 19, 20). Abolition by Δ*recA* can probably be explained by the need for homologous recombination both to establish the second “driver” of DnaA-independent replication as described above and to generate intact circular chromosomes from the newly replicated DNA regions; furthermore, any retrograde progression of the replisome (towards *oriC*) in these cells is likely to be impeded by frequent fork reversal events (89), whose resolution in Δ*recD* strains is known to be RecA-dependent (52, 88).

Leach and coworkers had used a strain in which SbcCD overexpression leads to cleavage of a palindrome on the lagging strand behind a replication fork to show that such cleavage leads in a Δ*recD* mutant to localized DNA amplification (64). In *recD*^*+*^ strains, the palindrome cleavage is expected to generate a one-ended DSB (since the *Ter*-proximal end would be resected back to the fork by RecBCD nuclease), whereas in a Δ*recD* mutant the consequence will be a two-ended DSB (106). Our model for ends-in replication outcomes in a Δ*recD* mutant would therefore account for their observations.

In summary, therefore, our proposal is that RecBCD’s ds-DNA end-resection activity is critically important to control rolling-circle over-replication at merger sites of converging replication forks in vivo. Models for chromosomal rolling-circle replication have also been proposed earlier in related but not identical contexts, by Kogoma and coworkers (99) for *recD* mutants (but not specifically at sites of fork merger) and by Marians and coworkers (107) for convergent replication forks in an in vitro system.

### Role of other DNA nucleases

Apart from RecJ, three other DNA nucleases Exo I, Exo VII, and SbcCD have been implicated in control of aberrant DnaA-independent chromosomal replication (37, 79, 108, 109); the quadruple mutant is cold-sensitive for growth and is UV^S^ (109, 110). We found that of these different DNA nucleases, it is RecJ deficiency alone that can singly synergize with Δ*recD* to rescue Δ*dnaA* lethality. It is likely that redundancy explains the failure of the other nucleases to do so. Rudolph, Lloyd, and coworkers have previously suggested that Exo I, Exo VII and SbcCD play redundant roles as 3’ ss-DNA exonucleases during the merger process of convergent replication forks in the terminus region (34, 37, 111), which may not be mutually exclusive with our model above.

## Data availability

The genome sequence and flow cytometry data described in this work are available for full public access from the repositories at https://www.ncbi.nlm.nih.gov/bioproject/842492 and https://flowrepository.org/id/FR-FCM-Z5F3, respectively.

## Supplementary data

Supplementary data are provided in a PDF file “Supplementary data”.

## Acknowledgements

We thank Olivier Espéli, Mohan Joshi, and Susan Rosenberg for strains; Nalini Raghunathan and Apuratha Pandiyan for assistance with flow cytometry and WGS data analysis; Manjula Reddy, Abhijit Sardesai, and R Harinarayanan for comments on the manuscript; and Christian Rudolph and COE team members for advice and discussions.

