## Supplementary for "Role for DNA double strand end-resection activity of RecBCD in control of aberrant chromosomal replication initiation in *Escherichia coli*"

**CONTENTS:**

**Supplementary text.**

**Supplementary References.**

**Table S1.** List of *E. coli* strains

**Table S2.** List of primer pairs for introduction of I-SceI sites on chromosome by recombineering

**Fig. S1.** Control experiments to elucidate roles of Δ*recD* mutation in rescue of Δ*dnaA* lethality.

**Fig. S2.** Impact of loss of RecA on viability of strains following different perturbations.

**Fig. S3.** Effects of *priA300* and Δ*dinG* on rescue of Δ*dnaA* lethality by Δ*recD* and Δ*recJ* mutations.

**Fig. S4.** Control flow cytometry experiments performed in absence of spectinomycin.

**Fig. S5.** Chromosomal copy number distributions, as determined by WGS, for **(A)** a control stationary phase culture of GJ18601 and **(B)** Δ*recJ* mutant GJ15990.

#### Supplementary text

**Recombineering of distributed I-SceI sites on chromosome.** With the exception of the site designated II (near *dusA*) the strain for which was obtained as gift from Mohan Joshi, the other four chromosomal I-SceI sites used in this study (I, III, IV, and V) were constructed by recombineering as follows. Plasmid pKD13 (1), that is commonly used as template for PCR in recombineering experiments, was first modified to carry an I-SceI site adjacent to the $Kan^R$ gene but outside the region bounded by the pair of FRT sequences. This was achieved (i) by the annealing of two complementary oligonucleotides (such as to leave 5'- end overhangs) 5'-TCGAGACTATAGGGATAACAGGGTAATAGCAC-3' and 5'-GATCGTGCTATTACCCTGTTATCCCTATAGTC-3', followed (ii) by ligation of the annealed duplex into the unique *SalI-BamHI* sites of pKD13 (2). The resultant plasmid pHYD5301 can be maintained in the S17-1 λ*pir* lysogenic strain.

In the second step, plasmid pHYD5301 was used as PCR template for recombineering of the I-SceI site along with FRT-flanked $Kan^R$ gene at different chromosomal locations; the primer pairs used for each location are listed in Table S2. Each recombineered I-SceI site was sequence verified. Derivatives with multiple I-SceI sites were constructed by serial successive steps of Flp recombinase-mediated excision of the $Kan^R$ element with the aid of plasmid pCP20 (1), followed by P1 transduction of the next construct.

**Table S1.** *List of E. coli strains*

| **Strain** [a] | **Genotype** [b] |
|---|---|
| **MG1655** | *E. coli* K-12 wild-type |
| **S17-1 $\lambda_{pir}$** | *recA1 endA1 thiE1 hsdR17* [$\lambda_{pir}$] |
| **GJ16464** [c] | GJ18601 Δ*dnaA*::FRT Δ*tus*::FRT *rpoB*35 btuB*::Tn*10* Δ*recG*::FRT |
| **GJ18601** | MG1655 Δ*(argF-lac)U169* |
| **GJ18604** [d] | GJ18601 Δ*dnaA*::FRT |
| **GJ18605** [d] | GJ18601 Δ*dnaA*::FRT Δ*tus*::FRT |
| **GJ18606** [d] | GJ18601 Δ*dnaA*::FRT *rpoB*35 btuB*::Tn*10* |
| **GJ18607** [d] | GJ18601 Δ*dnaA*::FRT Δ*tus*::FRT *rpoB*35 btuB*::Tn*10* |
| **GJ18609** | GJ18601 Δ*tus*::FRT *rpoB*35 btuB*::Tn*10* |
| **GJ15853** [d] | GJ18607 *dusA*::<(I-SceI$_{cutsite}$)-Sp> *att λ*::<*araC*(P$_{ara}$-I-SceI$_{enzyme}$)-Cm> |
| **GJ15857** [d] | GJ18607 *nadB*::<(I-SceI$_{cutsite}$)-FRT> *dusA*::<(I-SceI$_{cutsite}$)-Sp> *att λ*::<*araC*(P$_{ara}$-I-SceI$_{enzyme}$)-Cm> |
| **GJ15861** [d] | GJ18607 *nadB*::<(I-SceI$_{cutsite}$)-FRT> *dusA*::<(I-SceI$_{cutsite}$)-Sp> *yddT*::<(I-SceI$_{cutsite}$)-FRT> *att λ*::<*araC*(P$_{ara}$-I-SceI$_{enzyme}$)-Cm> |
| **GJ15864** [d] | GJ18607 *yddT*::<(I-SceI$_{cutsite}$)-FRT> *dusA*::<(I-SceI$_{cutsite}$)-Sp> *yaiZ*::<(I-SceI$_{cutsite}$)-FRT> *nadB*::<(I-SceI$_{cutsite}$)-FRT> *att λ*::<*araC*(P$_{ara}$-I-SceI$_{enzyme}$)-Cm> |
| **GJ15881** | S17-1$\lambda_{pir}$ (pHYD5301) |
| **GJ15888** [d] | GJ15853 Δ*recD*::Kan |
| **GJ15889** [d] | GJ18607 *yaiZ*::<(I-SceI$_{cutsite}$)-FRT> *dusA*::<(I-SceI$_{cutsite}$)-Sp> *att λ*::<*araC*(P$_{ara}$-I-SceI$_{enzyme}$)-Cm> Δ*recD*::Kan |
| **GJ15890** [d] | GJ18607 *yihQ*::<(I-SceI$_{cutsite}$)-FRT> *dusA*::<(I-SceI$_{cutsite}$)-Sp> *att λ*::<*araC*(P$_{ara}$-I-SceI$_{enzyme}$)-Cm> Δ*recD*::Kan |
| **GJ15892** [d] | GJ15857 Δ*recD*::Kan |
| **GJ15896** [d] | GJ15861 Δ*recD*::Kan |
| **GJ15899** [d] | GJ15864 Δ*recD*::Kan |
| **GJ15935** [d] | GJ18607 Δ*recD*::Kan |
| **GJ15936** [d] | GJ18607 Δ*recD*::FRT |
| **GJ15939** [d] | GJ18607 Δ*recD*::FRT Δ*rdgB*::Kan |
| **GJ15940** [d] | GJ18607 Δ*recD*::FRT *polA12 zih-3166*::Tn*10*Kan |
| **GJ15941** [d] | GJ18607 Δ*recD*::FRT Δ*dinG*::Kan |

| | |
|---|---|
| **GJ15942** [d] | GJ18607 Δ*recD*::FRT Δ*dinG*::Kan *att λ*::<($P_{ara}$-*rnhA*$^{+}$)-Amp> |
| **GJ15944** [d] | GJ18607 Δ*recD*::FRT *priA300* |
| **GJ15946** [d] | GJ18607 Δ*recD*::FRT Δ*xseA*::Kan |
| **GJ15947** [d] | GJ18607 Δ*recD*::FRT Δ*xonA*::Kan |
| **GJ15948** [d] | GJ18607 Δ*recD*::FRT Δ*sbcC*::Kan |
| **GJ15949** [d] | GJ18607 Δ*recD*::FRT Δ*recJ*::Kan |
| **GJ15970** [d] | GJ18607 Δ*dnaA*::Kan Δ*recD*::FRT Δ*recJ*::FRT Δ*dinG*::FRT |
| **GJ15971** [d] | GJ18607 Δ*dnaA*::Kan Δ*recD*::FRT Δ*recJ*::FRT Δ*dinG*::FRT *att λ*::<($P_{ara}$-*rnhA*$^{+}$)-Amp> |
| **GJ15973** [d] | GJ18607 Δ*dnaA*::Amp Δ*recD*::FRT Δ*recJ*::FRT *priA300* |
| **GJ15979** | MG1655 Δ*recD*::FRT Δ*recJ*::FRT |
| **GJ15981** | GJ18609 *rnhA339*::Cm |
| **GJ15984** | GJ18609 Δ*recD*::FRT |
| **GJ15985** | GJ18609 Δ*recJ*::FRT |
| **GJ15986** | GJ18609 Δ*recD*::FRT Δ*recJ*::FRT |
| **GJ15987** | GJ18609 Δ*recD*::FRT Δ*recJ*::FRT Δ*recA*::Kan |
| **GJ15988** | GJ18601 Δ*recD*::Kan |
| **GJ15990** | GJ18601 Δ*recJ*::Kan |
| **GJ15991** | GJ18601 Δ*recD*::FRT Δ*recJ*::Kan |
| **GJ15993** [d] | GJ18607 *tos*::Kan/N15 |
| **GJ15995** [d] | GJ18607 *rnhA339*::Cm *tos*::Kan/N15 |
| **GJ15999** [c] | GJ16464 *tos*::Kan/N15 |
| **GJ16001** [d] | GJ18607 Δ*recD*::FRT *tos*::Kan/N15 |
| **GJ16003** [d] | GJ18607 Δ*recD*::FRT Δ*recJ*::FRT *tos*::Kan/N15 |
| **GJ16072** [d] | GJ18604 Δ*recD*::Kan |
| **GJ16073** [d] | GJ18605 Δ*recD*::Kan |
| **GJ16078** [d] | GJ18606 Δ*recD*::Kan |
| **GJ16079** [d] | GJ18607 *att λ*::<*araC*($P_{ara}$-I-$SceI_{enzyme}$)-Cm> |
| **GJ16080** [d] | GJ18607 *att λ*::<*araC*($P_{ara}$-I-$SceI_{enzyme}$)-Cm> Δ*recD*::Kan |
| **GJ16081** [d] | GJ18607 *rnhA339*::Cm |
| **GJ16082** [d] | GJ18607 Δ*recD*::FRT *dut-1* |
| **GJ16083** [d] | GJ18607 Δ*recJ*::Kan |
| **GJ16084** [d] | GJ18607 Δ*recD*::FRT Δ*recJ*::FRT Δ*recA*::Kan |
| **GJ16088** [d] | GJ18607 Δ*dnaA*::Kan Δ*recD*::FRT Δ*recJ*::FRT Δ*dinG*::FRT *att λ*::<($P_{ara}$-Nil)-Amp> |
| **GJ16089** [d] | GJ18607 Δ*recD*::FRT Δ*dinG*::Kan *att λ*::<($P_{ara}$-Nil)-Amp> |

| | |
|---|---|
| **GJ16091** | GJ18609 $P_{sulA}$::(FRT-*lacZY*-Kan-*oriR6K*-FRT) |
| **GJ16092** | GJ18609 Δ*recD*::FRT $P_{sulA}$::(FRT-*lacZY*-Kan-*oriR6K*-FRT) |
| **GJ16093** | GJ18609 Δ*recA*::Kan |
| **GJ16094** | GJ18609 Δ*recD*::FRT Δ*recA*::Kan |
| **GJ16095** | GJ18609 Δ*recJ*::FRT Δ*recA*::Kan |
| **GJ16096** | GJ18609 Δ*recD*::FRT Δ*recJ*::FRT Δ*recA*::Kan |

[a] Strains MG1655, S17-1 $\lambda_{pir}$ , GJ18601, GJ18604, GJ18605, and GJ18606 were from our laboratory collection. Strains GJ16464, GJ18603, GJ18607, and GJ18609 have been described earlier (3). All other strains were constructed in this study.

[b] The following alleles and constructs have been described earlier: Δ(*argF-lac)U169* (4)*; btuB::*Tn*10* (3); Δ*dnaA*::Kan (3); *dut-1* (3); *polA12 zih-3166::Tn10*Kan (3) ; *priA300* (3); *rnhA339*::Cm (3); *rpoB*35* (3)*; att* λ::<*araC*($P_{ara}$-I-$SceI_{enzyme}$)-Cm> (5); *att* λ::<($P_{ara}$-Nil)-$Amp^r$> (3); *att* λ::<($P_{ara}$-*rnhA*$^+$)-Amp> (3); and $P_{sulA}$::(FRT-*lacZY*-Kan-*oriR6K*-FRT) (6). Prophage N15 and allele *tos*::Kan have also been described earlier (7), while the allele *dusA*::<*(I-SceI$_{cutsite}$)*-Sp> (designated by the notation II in Fig. 1A) was sourced from a strain provided by Mohan Joshi. The <(I-$SceI_{cutsite}$)-FRT> insertions at the following chromosomal locations (notations for each, as marked in Fig. 1A, are given in parentheses) *yihQ* (I), *yaiZ* (III), *yddT* (IV), and *nadB* (V) were generated in this study by recombineering, as described in the Supplementary text.

[c] Strain GJ16464 was routinely maintained as derivative with the *dnaA*$^+$ *recG*$^+$ shelter plasmid pHYD4805 (3).

[d] All other Δ*dnaA* strains were routinely maintained as derivatives with the *dnaA*$^+$ shelter plasmid pHYD2388 (3).

**Table S2.** *List of primer pairs for introduction of I-SceI sites on chromosome by recombineering*

| Chromosomal locus | Notation in Fig. 1A | Primer sequence pairs (5'-3') |
|---|---|---|
| *yihQ* | **I** | TAATTCATTTCAGCCGTGGTTCTGACATTAGCGCCACGCTG ATTGTGTAGGCTGGAGCTG |
| | | TTCCAGCAATAAACGCCCCTGATCGTCGGCAGAGATATTCC ACTGATCAGTGATAAGCTG |
| *yaiZ* | **III** | ATATTCATTCTTAATGGCGTGGTGGGGTTACTGGGATTTGG ATTGTGTAGGCTGGAGCTG |
| | | GTCACCAGACCGACGGCATAGGTCTGCCAACCTTTTGCTTC ACTGATCAGTGATAAGCTG |
| *yddT* | **IV** | ACAAGGATGCGATTACCGCGCTGGCGAAAAGTATCAGCAT GATTGTGTAGGCTGGAGCTG |
| | | CGCATCGTAAATCACCAGTTGTAACCCTGACAGCTGGGCGC ACTGATCAGTGATAAGCTG |
| *nadB* | **V** | GTAACGGAAGGTTCAACATTTTATGCCCAGGGCGGTATTGG ATTGTGTAGGCTGGAGCTG |
| | | CACATGCGAGTCAATGCTGTCAGTTTCATCAAACACGGCGC ACTGATCAGTGATAAGCTG |

**Figure S1**

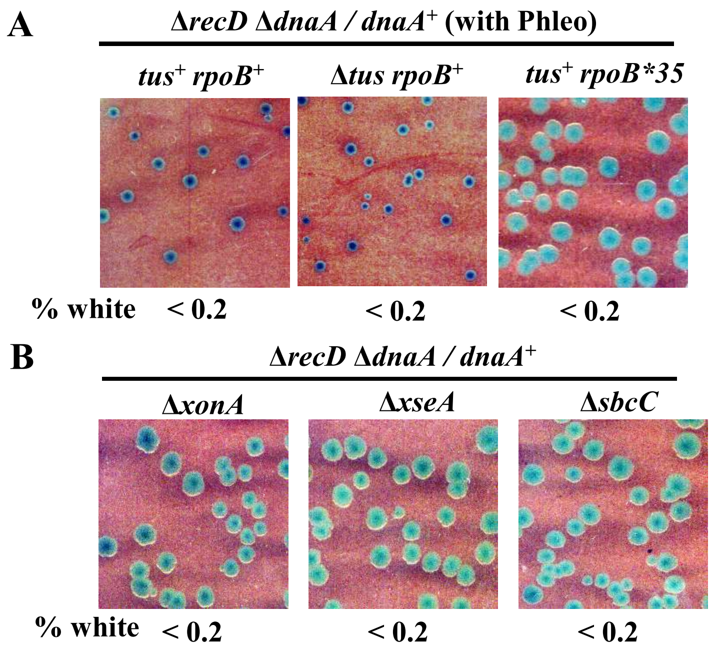


**Figure S1**: Control experiments to elucidate roles of Δ*recD* mutation in rescue of Δ*dnaA* lethality. Blue-white assay results are depicted, as described in legend to Fig. 1B, for **(A)** Δ*recD* derivatives with different combinations of *tus* and *rpoB* genotypes, plated on LB supplemented with Phleo; and **(B)** Δ*recD* derivatives bearing the different additional single mutations as indicated on top of each sub-panel. Strains used for the different sub-panels were (from left, all strain numbers are prefixed with GJ): A, 16072, 16073, and 16078; and B, 15947, 15946, and 15948.

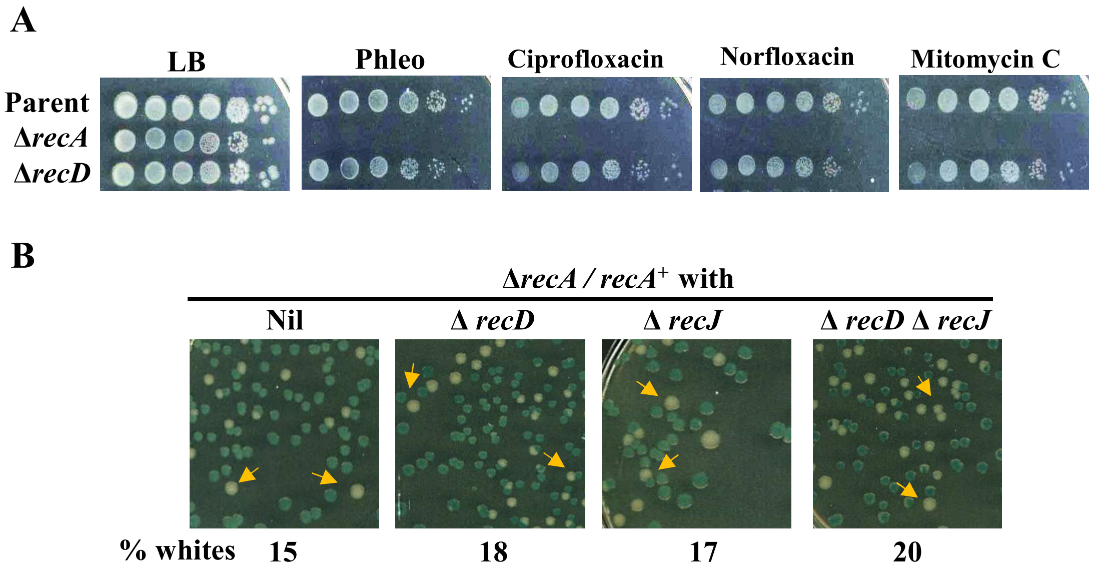


**Figure S2**: Impact of loss of RecA on viability of strains following different perturbations. **(A)** Dilution-spotting assays on medium without or with supplementation with genotoxic agents as indicated. Concentrations employed were the same as those described in the text and in legend to Fig. 3A. Strains used were (all strain numbers are prefixed with GJ): Parent, 18609; Δ*recA*, 16093; and Δ*recD*, 15984. **(B)** Blue-white assay results are depicted, as described in legend to Fig. 1B, for different chromosomal Δ*recA* derivatives bearing *recA*$^{+}$ shelter plasmid pHYD5701. Strains used were (all strain numbers are prefixed with GJ): Nil, 16093; Δ*recD*, 16094; Δ*recJ*, 16095; and Δ*recD* Δ*recJ*, 16096.

### Figure S3

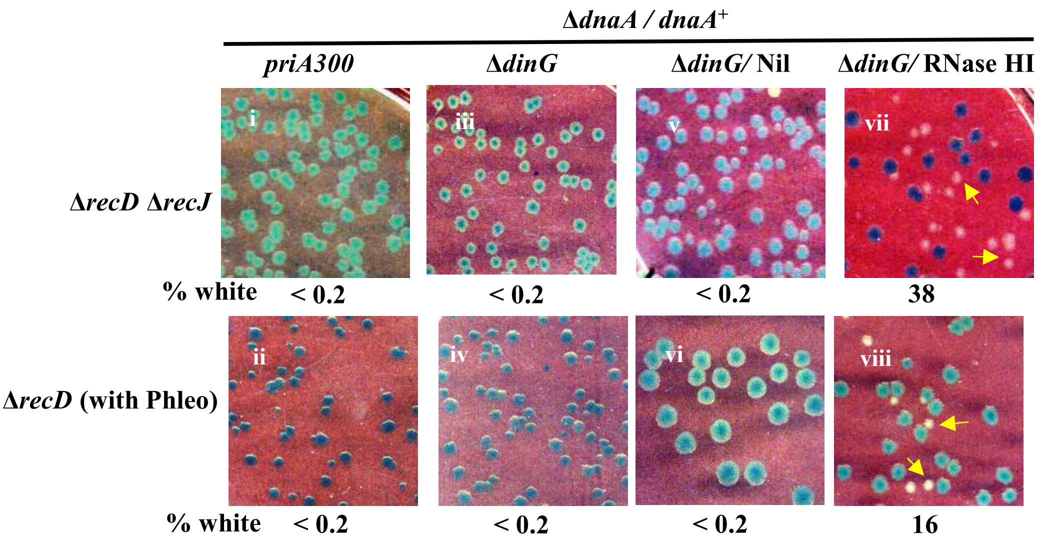


**Figure S3**: Effects of *priA300* and Δ*dinG* on rescue of Δ*dnaA* lethality by Δ*recD* and Δ*recJ* mutations. Blue-white assay results are depicted, as described in legend to Fig. 1B, for Δ*recD* Δ*recJ* derivatives (top row) and Δ*recD* derivatives plated on LB with Phleo (bottom row), with additional genetic perturbations as indicated on top of each pair of panels. Strains used for the different panels were (all strain numbers are prefixed with GJ): i, 15973; ii, 15944; iii, 15970; iv, 15941; v, 16088; vi, 16089; vii, 15971; and viii, 15942.

**Figure S4**

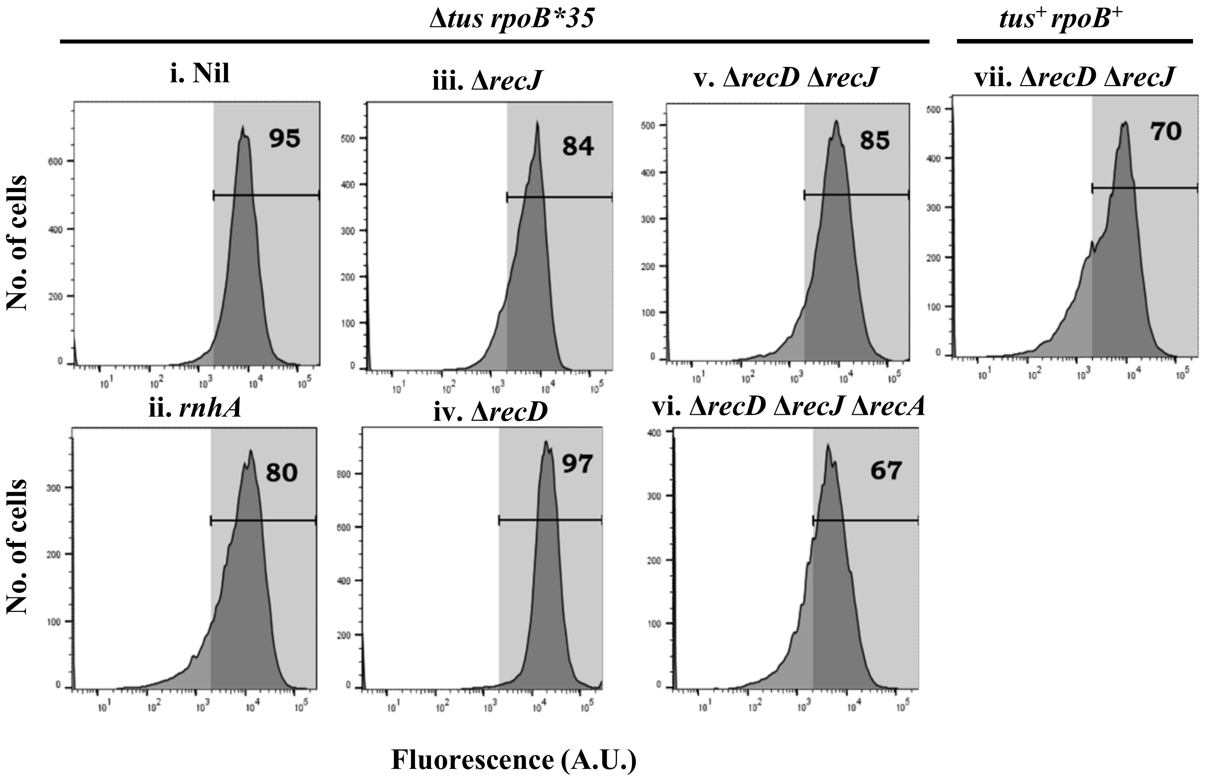


**Figure S4**: Control flow cytometry experiments performed in absence of spectinomycin. The depicted panels are the paired controls of those in Fig. 5, with the difference that these experiments were performed without addition of spectinomycin to the cultures.

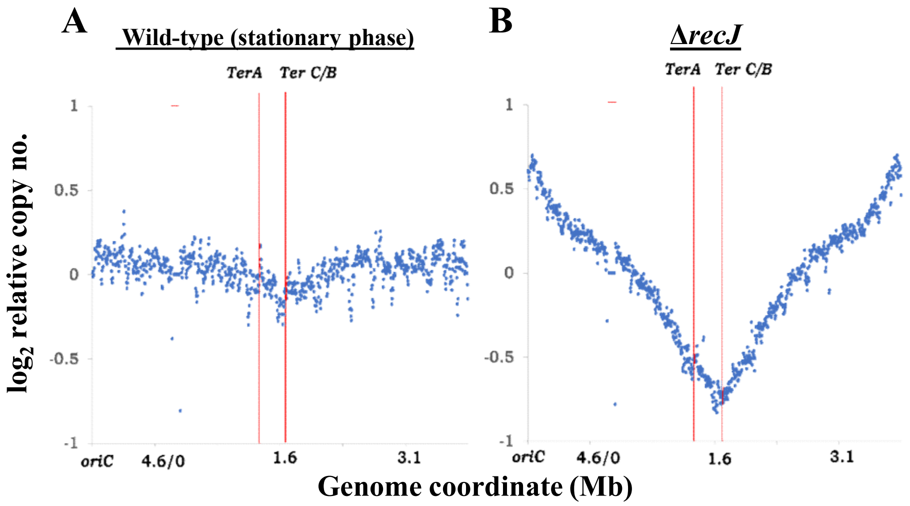


**Figure S5**: Chromosomal copy number distributions, as determined by WGS, for **(A)** a control stationary phase culture of wild-type strain GJ18601 and **(B)** Δ*recJ* mutant GJ15990. Representations of WGS analysis data and notations used are as described in the legend to Fig. 6.
